# Hierarchical chromatin polyvalency governs robust gene regulation and organogenesis

**DOI:** 10.64898/2026.09.15.751898

**Authors:** Chengjie Zhou, Meng Wang, Zhiyuan Chen, Yi Zhang

**Affiliations:** Howard Hughes Medical Institute, Boston Children’s Hospital, Boston, MA 02115, USA; Program in Cellular and Molecular Medicine, Boston Children’s Hospital, Boston, MA 02115, USA; Reproductive Sciences Center, Division of Developmental Biology, Cincinnati Children’s Hospital Medical Center, Cincinnati, OH 45229, USA; Department of Pediatrics, University of Cincinnati College of Medicine, Cincinnati, OH 45267, USA; Division of Hematology/Oncology, Department of Pediatrics, Boston Children’s Hospital, Boston, MA 02115, USA; Department of Genetics, Harvard Medical School, Boston, MA 02115, USA; Harvard Stem Cell Institute, Boston, MA 02115, USA

**Keywords:** Polyvalency, H2Aub, H3K27me3, H3K4me3, H3K9me3, organogenesis, somitogenesis

## Abstract

Precise temporal control of gene expression is fundamental for embryonic development, yet the epigenetic and chromatin basis governing transcriptional timing remain poorly understood. The bivalency model, characterized by coexistence of H3K4me3 and H3K27me3, has been proposed to mark a poised state ready for activation. However, this model has been challenged by lacking of rapid gene activation in response to H3K27me3 depletion, suggesting that the H3K27me3 mark is not responsible for the silencing. Here, through temporal epigenomic profiling of post-implantation mouse embryos and use of the protein degradation tag (dTAG) system, we demonstrate that H2Aub, but not H3K27me3, functions as the major repressor. We further reveal a hierarchical repression architecture in which H2Aub is responsible for transcriptional silencing, while H3K27me3 and H3K9me3 serve to reinforce the silencing state in post-implantation embryos. Functionally, disruption of this H2Aub-centered hierarchy perturbs temporal control of polyvalent gene activation, leading to severe organogenesis defects. Mechanistically, acute loss of H2Aub disrupts retinoic acid-FGF signaling pathway, causing somitogenesis arrest. Together, our findings establish chromatin polyvalency model as a multi-layered, hierarchical repression mechanism that governs temporal control of gene expression during embryogenesis.

## Introduction

Epigenetic modifications play a fundamental role in regulating gene activation and silencing through a set of key histone and DNA modifications ^1–3^. Among the modifications, SET1/COMPASS family enzymes-catalyzed histone H3 lysine 4 trimethylation (H3K4me3) facilitates gene activation by promoting paused RNA polymerase II (Pol II) release ^4^. While the Polycomb Repressive Complex 1 (PRC1) deposited histone H2A lysine 119 mono-ubiquitination (H2Aub) and PRC2 catalyzed histone H3 lysine 27 trimethylation (H3K27me3) mediate gene repression by inhibiting RNA Pol II recruitment and elongation^5–12^. PRC1 complexes are broadly classified into canonical (cPRC1) and variant (vPRC1) subtypes based on their subunit composition. cPRC1 contains PCGF2/4 and a CBX chromodomain-containing subunit that recognizes H3K27me3, whereas vPRC1 incorporates one of six PCGF paralogs together with RYBP/YAF2 and mediates the majority of H2AK119ub1 deposition in an H3K27me3-independent manner ^13, 14^. In contrast to Polycomb silencing, histone H3 lysine 9 trimethylation (H3K9me3) mediates long-term gene silencing by establishing constitutive heterochromatin ^15, 16^. Together, these histone markers constitute the basic framework of our current understanding of epigenetic regulation of gene expression.

Precise temporal control of gene expression is essential for diverse biological processes, particularly those involving rapid cell state transitions, such as immune activation, stress responses and embryonic development ^17, 18^. These processes require swift and tightly regulated gene activation and silencing to ensure appropriate cellular responses ^19, 20^. In such contexts, a group of key regulatory genes are transcriptionally repressed or maintained at low expression levels, yet remain poised for rapid activation upon receipt of appropriate signals ^21–23^. This poised state enables cells to balance stability with plasticity ^24^. However, although the importance of transcriptional poising is well established, the chromatin-basis that maintain such state while preventing premature gene activation is not fully understood. Among the proposed models, chromatin bivalency has emerged as a widely accepted framework for explaining how genes can remain repressed yet poised for rapid activation ^21, 23^. The classic bivalency is defined by the coexistence of the activating marker H3K4me3 and the repressive marker H3K27me3 ^21^. Bivalent chromatin was first identified at developmental genes in pluripotent ESCs, and later confirmed in many other cell types, including neural progenitor cells, cancer cells and post-implantation embryos ^25–29^. In this model, H3K27me3-mediated repression is thought to counteract transcriptional activation and maintain genes in a silent but poised state. Rapid removal of H3K27me3 was supposed to resolve the bivalent chromatin configuration, leading to rapid transcriptional activation of H3K4me3-marked promoters to ensure timely gene activation during embryonic development. However, genetic and functional studies have shown that acute H3K27me3 depletion does not necessarily lead to transcriptional activation of bivalent genes ^30^, suggesting that the current bivalency model needs to be redefined in order to reconcile the observations. Previous studies have shown that H2Aub is enriched at a subset of bivalent genes in mouse ES cells ^31, 32^, suggesting that, in addition to H3K27me3, other histone modification may also contribute to the regulation of bivalency. However, how these histone modifications cooperate to maintain bivalent genes in a poised yet repressed state remains poorly understood. More importantly, it remains unclear whether insights derived from stable cultured cells can be applied to the highly dynamic and extensive cell lineage differentiation that occurs during post-implantation embryonic development.

To address these questions, we conducted temporal epigenomic profiling of histone modifications around mouse gastrulation stages, a developmental window characterized by rapid developmental progression and simultaneous specification of multiple cell lineages ^33, 34^, making the embryonic cells an ideal context for investigating transcriptional poising and the precise, reversible regulation of gene expression. Here we identify a polyvalent chromatin state in which multiple histone modifications (H3K4me3/H3K27me3/H2Aub/H3K9me3) coexist at gene promoters *in vivo*. By using a protein degradation tag (dTAG) approach ^35–37^, we acutely depleted PRC1 or PRC2 in a developmental stage-specific manner, and revealed that PRC1, but not PRC2, is the key repressor of bivalent genes. Our findings reveal an H2Aub-centered regulatory network involving H3K27me3, H3K9me3 and H3K4me3 that collectively constitutes the polyvalent chromatin state during post-implantation development. Importantly, the use of the dTAG system allowed us to discover a critical role of PRC1 in regulating somitogenesis by modulating FGF signaling and wavefront through polyvalent genes, which is not possible by conventional genetic approach due to the 2-cell or 4-cell arrest phenotype of PRC1 depletion ^38, 39^. These results also highlight a previously unknown molecular mechanism by which PRC1 regulates somitogenesis through RA/FGF signaling.

## Results

### Acute PRC2/H3K27me3 degradation has minor effects on bivalent gene expression

A comprehensive analysis of H3K27me3 and H3K4me3 CUT&RUN dataset identified thousands of bivalent genes in mouse embryonic day 6.5 (E6.5) epiblast (EPI), extraembryonic ectoderm (ExE), visceral endoderm (VE), and embryonic stem cells (ESCs), respectively (**Fig. 1a, b and Supplementary Table 1**). To determine whether acute loss of PRC2 could activate these bivalent genes, we used the *Eed* dTAG mouse model described in our previous study (**Fig. 1c**) ^40^. This system enables rapid degradation of EED and removal of H3K27me3 within 12 hours following *in vivo* dTAG^V^-1 injection (**Extended Data Fig. 1a**). Surprisingly, acute EED depletion from E5.5 to E6.5 had minimal effects on bivalent gene expression in EPI (1 of 2,606, 0.04%) and VE (2 of 1,944, 0.10%) lineage, and only 87 bivalent genes (4.44%) were upregulated in ExE (**Fig. 1d and Supplementary Table 2**). Although the bivalent genes showed a trend toward upregulation in ExE lineage, the majority of the bivalent genes remained silent (**Fig. 1e**). Similar to bivalent genes, the H3K27me3-only genes in all three lineages showed minor changes upon H3K27me3 depletion (**Extended Data Fig. 1b**).

**Figure 1.**
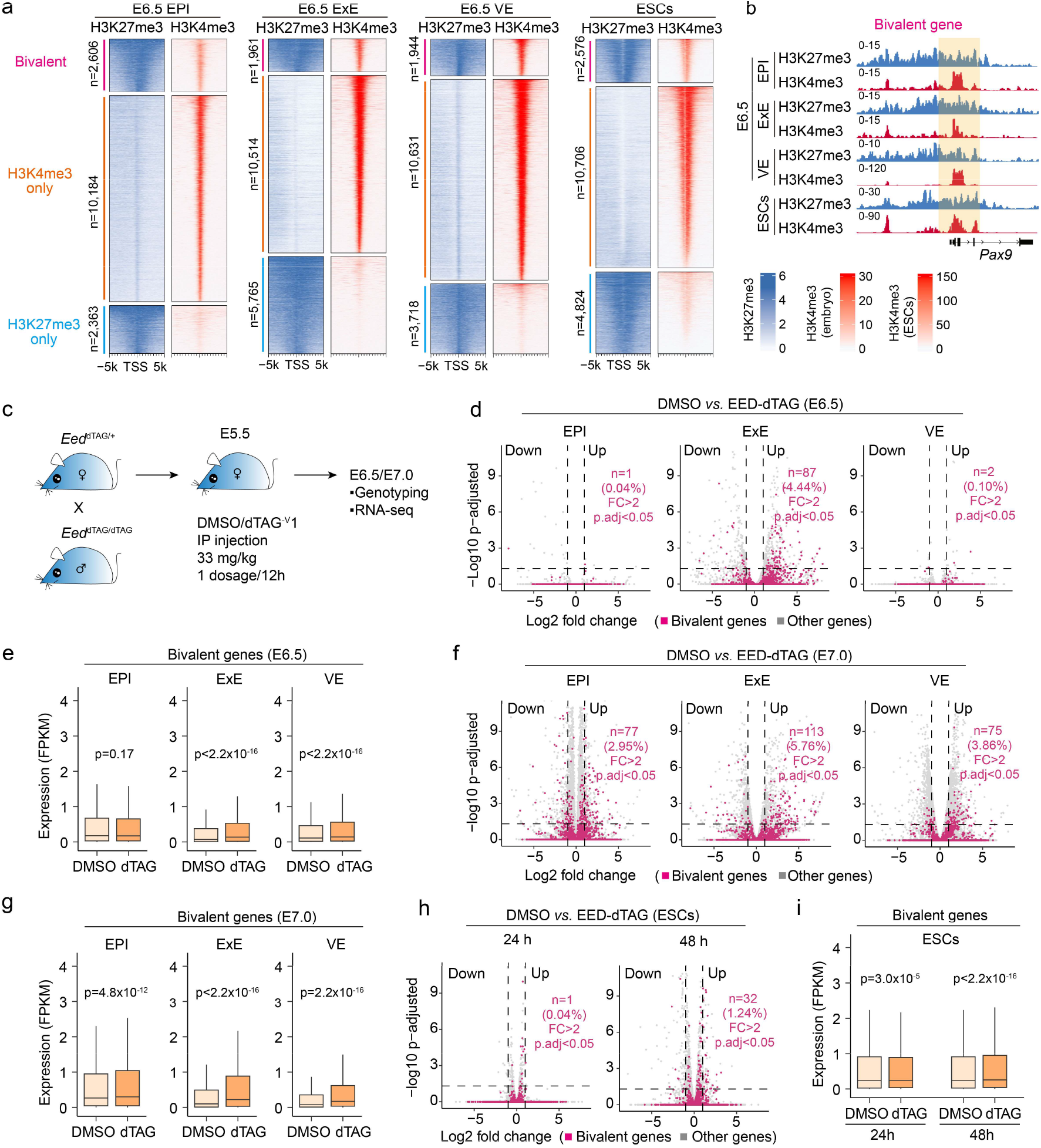
Acute PRC2 depletion has minor effects on bivalent genes expression. (**a**) Heatmaps showing the gene classification based on the H3K27me3 and H3K4me3 levels around the transcription start sites (TSS) in E6.5 EPI, ExE, VE and ESCs. (**b**) Genome browser views of H3K27me3 and H3K4me3 at the *Pax9* gene locus in E6.5 EPI, ExE, VE and ESCs. (**c**) Schematic showing the experimental design for dTAG^V^-1 mediated EED degradation. (**d**) Volcano plots comparing gene expression changes between DMSO and EED-depleted E6.5 EPI, ExE and VE cells. Red dots represent bivalent genes, and gray dots represent non-bivalent genes. FC: fold change. (**e**) Box plot showing the RNA expression changes of bivalent genes in DMSO and EED-depleted E6.5 EPI, ExE and VE cells. p values were calculated with Wilcoxon signed-rank test (two-tailed). (**f**) Volcano plots comparing gene expression between DMSO and EED-depleted E7.0 EPI, ExE and VE cells. Red dots represent bivalent genes. (**g**) Box plots showing the RNA expression changes of bivalent genes in DMSO and EED-depleted E7.0 EPI, ExE and VE cells. p values were calculated with Wilcoxon signed-rank test (two-tailed). (**h**) Volcano plots comparing gene expression changes between DMSO and EED-depleted ESCs at 24 and 48 hrs of dTAG treatment. (**i**) Box plots showing the RNA expression changes of bivalent genes in DMSO and EED-depleted ESCs. p values were calculated with Wilcoxon signed-rank test (two-tailed). For all the boxplots, the central band represents the median. The lower and upper edges of the box represent the first and third quartiles, respectively. The whiskers of the boxplot extend to 1.5 times interquartile range (IQR).

To assess whether prolonged EED depletion increases bivalent gene activation, we performed RNA-seq analyses at E7.0 following EED degradation (**Extended Data Fig. 1c**). Compared with E6.5, E7.0 EPI, ExE and VE showed more differentially expressed genes (**Extended Data Fig. 1d**) with more bivalent genes upregulated (EPI, 2.95%; ExE, 5.76%; VE, 3.86%) after additional 12-hour EED degradation (**Fig. 1f, g and Supplementary Table 2**). Nevertheless, the great majority of bivalent genes still remained repressed. To test whether acute PRC2 loss affects bivalent genes expression in ESCs, we induced complete EED degradation by treating ESCs with dTAG-13 for 24 or 48 hours (**Extended Data Fig. 1e**). Consistent with the *in vivo* results, few bivalent genes were up-regulated at 24 (1 out of 2,576, 0.04%) and 48 hours (32 out 2,576 of 1.24%) (**Fig. 1h, Extended Data Fig. 1f and Supplementary Table 2**), although an overall upward trend was observed (**Fig. 1i**). These results demonstrate that PRC2 and H3K27me3 removal is not sufficient to activate bivalent genes *in vivo* and *in vitro*.

Collectively, these findings indicate that although bivalent genes are classically defined by the coexistence of H3K27me3 and H3K4me3, acute PRC2/H3K27me3 depletion only has a minor effect on their expression. This suggests that the traditional H3K27me3/H3K4me3 bivalency model is largely descriptive rather than functional, which raises the possibility that additional repressive mechanisms are responsible for maintaining the silencing of these bivalent genes.

### H2Aub and H3K9me3 are present on bivalent genes in post-implantation embryos

Several transcription repressive histone markers, including H3K27me3, H2Aub, and H3K9me3, have been identified ^41^. Among these repressive markers, H3K27me3 and H2Aub respectively deposited by PRC2 and PRC1 usually colocalized ^5, 7, 8^. In addition, H3K4me3 and H3K9me3 can also form a bivalent state ^42, 43^. These finding suggest that H2Aub and H3K9me3 might be involved in bivalent gene silencing. To test this possibility, we asked whether H2Aub and H3K9me3 are associated with bivalent genes by performing CUT&RUN profiling for H2Aub and H3K9me3 in E6.5 EPI, ExE and VE lineages (**Extended Data Fig. 2a**). This analysis revealed that bivalent genes (cluster 1, C1) in EPI and ESCs were highly enriched for both H2Aub and H3K9me3, whereas H2Aub was less enriched in the ExE lineage and H3K9me3 was less enriched in the VE lineage (**Fig. 2a, b**). We next subdivided bivalent genes into four groups based on their combinatorial histone modification states (**Fig. 2c, d**). Interestingly, the majority of the bivalent genes enriched with both H2Aub and H3K9me3 (EPI: 62.97%, ExE: 78.23%, VE: 30.76%) or H2Aub (K27/K4/H2A, EPI: 37.03%, ExE: 21.11%, VE, 69.19%) (**Fig. 2c**). In contrast, very few bivalent genes were marked only by H3K27me3 and H3K4me3. A pattern similar to that of the EPI and ExE lineages is observed in ESCs (**Fig. 2c**). Together, these results indicate that genes previously classified as bivalent are not only marked by H3K4me3 and H3K27me3, but harbor additional repressive markers, including H2Aub and H3K9me3.

**Figure 2.**
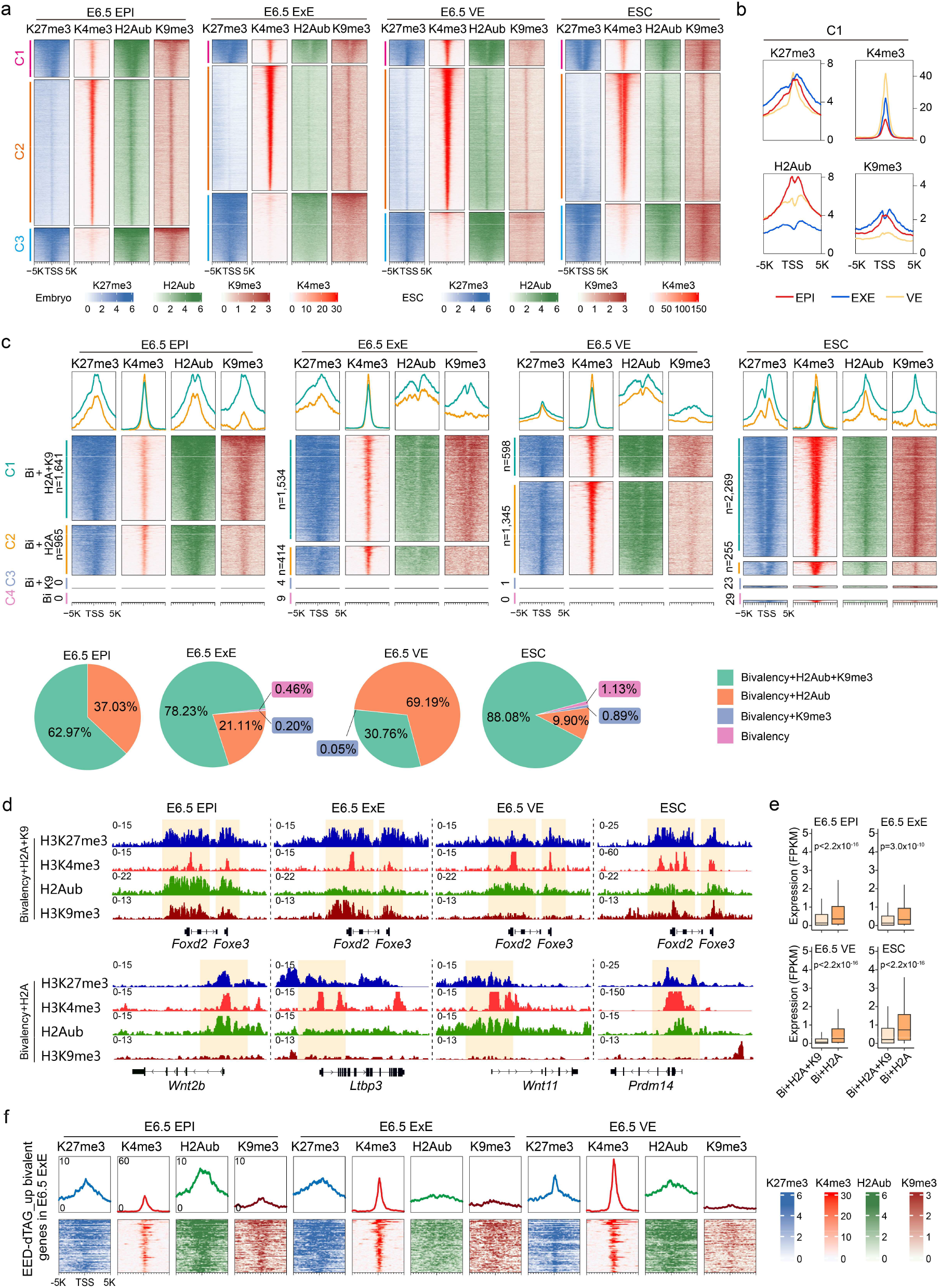
H2Aub and H3K9me3 are also present on bivalent genes. **(a)** Histone modification profiling reveals that bivalent genes (cluster 1, C1) also harbor H2Aub and H3K9me3 modifications in E6.5 EPI, ExE, VE and ESCs. (**b**) Meta plots showing the H3K27me3, H3K4me3, H2Aub and H3K9me3 levels in the bivalent genes (C1) in E6.5 EPI, ExE and VE cells. (**c**) Histone modification profiling identifies a set of bivalent genes enriched with H2Aub or H3K9me3. The pie charts show the percentage of genes in each category. (**d**) Genome browser views of bivalent gene examples enriched with H2Aub or H3K9me3 in E6.5 EPI, ExE, and VE cells. (**e**) Box plots showing expression of bivalent genes with different histone markers in E6.5 EPI, ExE and VE, as well as ESCs. In the boxplot, the central band represents the median. The lower and upper edges of the box represent the first and third quartiles, respectively. The whiskers of the boxplot extend to 1.5 times interquartile range (IQR). p values were calculated with two-sided Mann-Whitney U test. (**f**) Heatmaps and meta plots showing enrichment of H3K27me3, H3K4me3, H2Aub and H3K9me3 on bivalent genes that showed upregulation in E6.5 ExE (n=87).

While these findings expand the classical definition of bivalency, the role of these additional marks in regulating bivalent gene expression remains to be shown. Consistent with the higher levels of H2Aub and H3K9me3 observed at genes of Cluster 1 (H3K27me3+H3K4me3+H2Aub+H3K9me3) relative to the genes of Cluster 2 (H3K27me3+H3K4me3+H2Aub) (**Fig. 2c**), bivalent genes enriched with H2Aub alone exhibited significantly higher expression than that enriched with both H2Aub and H3K9me3 in EPI, ExE, VE and ESCs (**Fig. 2e**), suggesting that H3K9me3 may contributes to the repressive state of these bivalent genes. We next asked whether lineage-specific differences in H2Aub and H3K9me3 levels could explain the selective upregulation of bivalent genes in the ExE, but not in the EPI or VE, following acute EED degradation (**Fig. 1d, f**). To this end, we analyzed epigenetic features of bivalent genes upregulated in E6.5/E7.0 ExE following acute EED degradation. Compared with ExE, EPI exhibited comparable levels of H3K27me3 and H3K9me3 but much higher levels of H2Aub, whereas VE exhibited higher H2Aub and lower H3K9me3 levels (**Fig. 2f and Extended Data Fig. 2b**). These results suggest that H2Aub, rather than H3K9me3, may function as a functional repressive mark in maintaining the silencing state in both EPI and VE when PRC2-mediated H3K27me3 is removed.

### Acute PRC1 degradation activates bivalent genes

We next sought to determine whether depletion of the H2Aub mark through rapid degradation of the core PRC1 components could activate bivalent genes. To this end, we generated the *Ring1a* and *Ring1b* dTAG mice by inserting the *Fkbp12^F36V^-HA* degron before the stop codon of the endogenous *Ring1a* locus and after the start codon of the endogenous *Ring1b* locus with CRISPR technology, respectively (**Extended Data Fig. 3a-c**). The *Ring1b*^dTAG/dTAG^ homozygous were viable and exhibit normal litter sizes (**Extended Data Fig. 3d**). *Ring1a*^dTAG/dTAG^; *Ring1b*^dTAG/dTAG^ double*-*homozygous showed normal development from gastrulation to late gestation (**Extended Data Fig. 3e**), indicating that combined *Ring1a*-dTAG and *Ring1b*-dTAG does not impair embryogenesis. These embryos therefore provide a suitable and robust model for investigating early post-implantation development (**Extended Data Fig. 3e**).

To evaluate RING1A/1B and H2Aub degradation efficiency, *Ring1a*^dTAG/dTAG^; *Ring1b*^dTAG/+^ female mice were used to mate with *Ring1a*^dTAG/dTAG^*; Ring1b*^dTAG/+^ male mice, and subjected to intraperitoneal injection of dTAG^V^-1 at E5.5 (**Fig. 3a**). Immunostaining of E6.5 embryos confirmed that RING1A/1B and H2Aub were completely depleted in both male and female embryos after ∼24-hr dTAG^V^-1 treatment (**Fig. 3b**). We then dissected E6.5 EPI, ExE and VE for RNA-seq analyses (**Extended Data Fig. 4a**). Strikingly, acute RING1A/1B degradation led to 151 (5.83%) and 643 (24.67%) bivalent genes upregulation (FC≥2, adjusted p<0.05) in E6.5 EPI and E7.0 EPI, respectively (**Fig. 3c and Supplementary Table 3**). Interestingly, bivalent genes showed a strong bias toward the upregulation on the volcano plot (**Fig. 3c)**, while the active genes (cluster 2 in Fig. 2a) did not or only show a weak bias (**Extended Data Fig. 4b**). Consistent with this observation, the global bivalent genes showed significant upregulation upon acute RING1A/1B depletion in both E6.5 and E7.0 EPI (**Fig. 3d**). Moreover, PRC1 also displayed a strong repressive effect on bivalent genes in VE and ESCs, but not in ExE (**Fig. 3e, f and Extended Data Fig. 4c-j**), which is consistent with the lower H2Aub intensity in ExE lineage, compared with other lineages (**Fig. 2a, b**).

**Figure 3.**
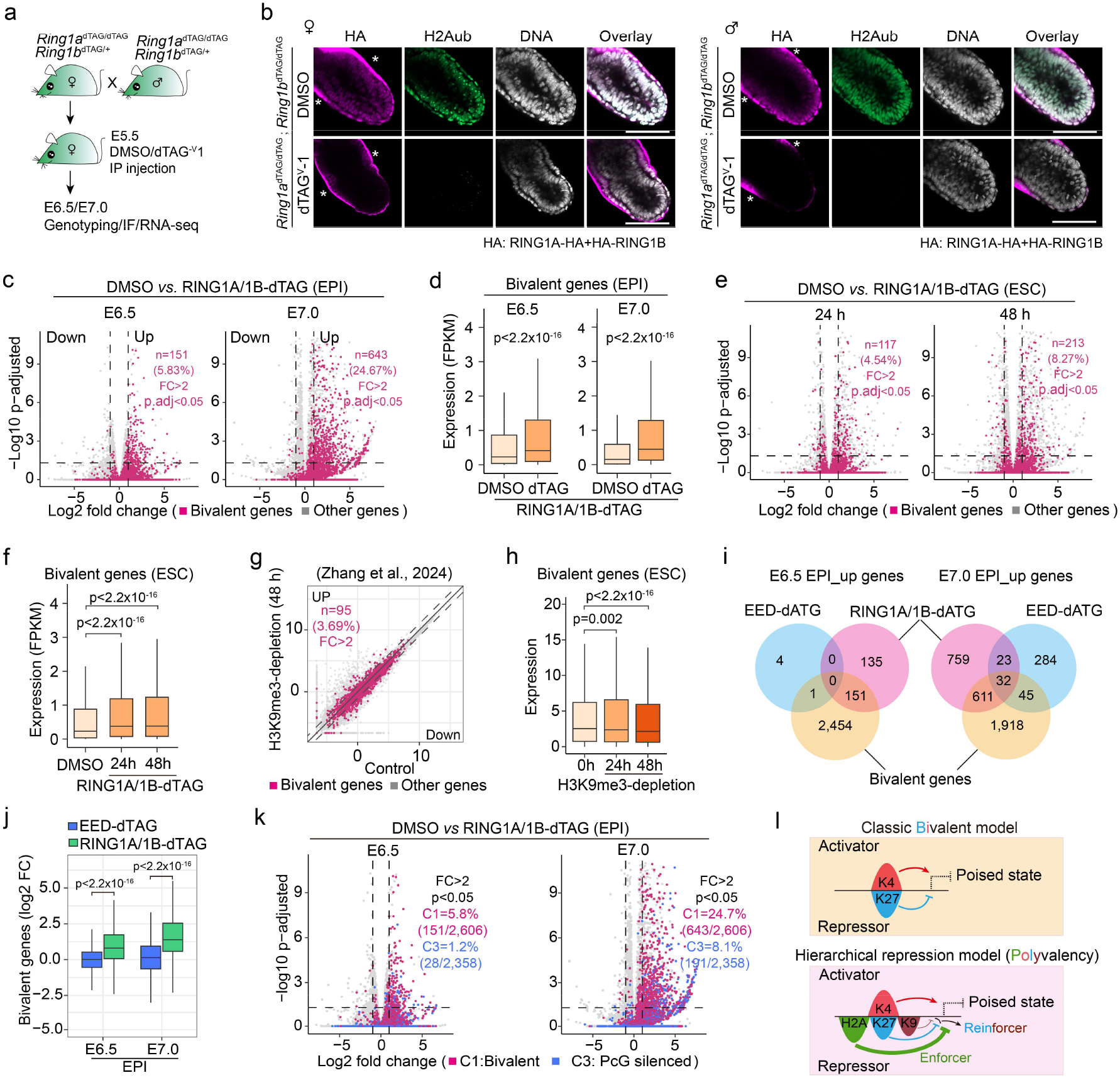
PRC1, but not PRC2, plays a major role in silencing bivalent genes. (**a**) Schematic illustrating the experimental design for dTAG^V^-1 mediated RING1A and RING1B degradation *in vivo*. (**b**) Immunostaining showing the rapid depletion of RING1A, RING1B and H2Aub in E6.5 embryos (female and male) following intraperitoneal (IP) injection of dTAG^V^-1. All embryos were collected, genotyped, and processed for downstream analyses. HA staining indicates expression of HA-RING1A and RING1B-HA. Asterisks denote non-specific staining. DNA was stained with Hoechst 33342. Scale bar, 100 μm. (**c**) Volcano plots comparing gene expression changes between DMSO and RING1A/1B-depleted E6.5 and E7.0 EPI. Red dots represent bivalent genes (n=2,606), and gray dots represent non-bivalent genes. FC: fold change. (**d**) Box plot showing the RNA expression changes of bivalent genes in DMSO and RING1A/1B-depleted E6.5 and E7.0 EPI. p values were calculated with Wilcoxon signed-rank test (two-tailed). (**e**) Volcano plots comparing gene expression changes following 24 h and 48 h dTAG treatment for RING1A/1B depletion in ESCs. Red dots represent bivalent genes (n=2,576), and gray dots represent non-bivalent genes. (**f**) Box plots showing expression changes of bivalent genes in ESCs following 24 h and 48 h dTAG treatment for RING1A/1B depletion. p values were calculated with Wilcoxon signed-rank test (two-tailed). (**g**) Transient transcriptome sequencing (TT-seq) comparison of ESCs with or without H3K9me3 depletion for 48 h (Zhang et al., 2024). (**h**) Box plot showing the comparison of bivalent gene expression with or without H3K9me3 depletion for 24 and 48 h in ESCs. p values were calculated with Wilcoxon signed-rank test (two-tailed). (**i**) Venn diagram showing the overlap of upregulated genes upon EED or RING1A/1B depletion and their overlap with bivalent genes in E6.5 and E7.0 EPI. (**j**) Comparison of log₂ fold changes of bivalent genes following EED or RING1A/1B depletion in E6.5 and E7.0 EPI. p values were calculated with Wilcoxon signed-rank test (two-tailed). (**k**) Volcano plots showing differential gene expression in E6.5 and E7.0 EPI upon RING1A/1B-dTAG treatment compared with DMSO. Red dots represent bivalent genes (C1 in Fig. 2a), blue dots represent PcG-silenced genes (C3 in Fig. 2a). (**l**) Schematic model of polyvalent chromatin, highlighting the hierarchical repression by H2Aub, H3K27me3, and H3K9me3 to maintains a poised chromatin state. K4, H3K4me3; H2A, H2Aub; K27, H3K27me3; K9, H3K9me3. For all the boxplots, the central band represents the median. The lower and upper edges of the box represent the first and third quartiles, respectively. The whiskers of the boxplot extend to 1.5 times interquartile range (IQR).

We next sought to investigate whether H3K9me3 contributes to the silencing of bivalent genes. To this end, we analyzed public nascent transcriptional profiles ^44^ of ESCs comparing control with H3K9me3 depletion. We found that only a small group of bivalent genes were upregulated (n=95 at 24 h and n=81 at 48 h after H3K9me3 depletion, FC≥2) (**Fig. 3g and Extended Data Fig. 4k**), while global expression of bivalent genes remained largely unchanged (24 h) or modestly reduced (48 h), potentially due to indirect effects (**Fig. 3h**). These results are consistent with *in vivo* observation that the presence or absence of H3K9me3 does not alter PRC1’s role in silencing bivalent genes (**Extended Data Fig. 4l, m**). Collectively, these analyses demonstrate that acute loss of H2Aub, but not H3K27me3 or H3K9me3, results in a broad derepression of bivalent genes, supporting H2Aub serves as the central repressor in bivalent gene silencing. This finding underscores a dominant and indispensable role of H2Aub in maintaining the silencing state of bivalent genes.

Although H3K27me3 has been regarded as the repressive component in classic bivalent model, acute RING1A/1B degradation resulted in upregulation of more genes and stronger transcriptional activation than EED degradation (**Fig. 3i, j and Extended Data Fig. 4n, o**), indicating that H2Aub serves as a more potent repressive mark than H3K27me3 in this context. Notably, loss of PRC1 activates more bivalent genes (C1 in **Fig. 2a**) than the PcG silenced genes (C3 in **Fig. 2a**) (**Fig. 3k)**, suggesting that loss of H2Aub is the primary trigger for releasing the poised state during bivalent gene activation. Given that EED/H3K27me3 depletion only modestly activates bivalent genes, the cPRC1/H3K27me3 is unlikely to account for the transcriptional activation upon RING1A/B loss. These results suggest that vPRC1-mediated H2Aub should account for the major PRC1 function maintaining bivalent genes in a poised, repressed state. Together, these results indicate that the classic bivalent model primarily describes co-occupancy of H3K27me3 and H3K4me3 rather than providing a functional explanation of gene regulation. To address this limitation, we propose a hierarchical repression model (named polyvalency) that describes a chromatin state with co-occurrence of three repressive makers H2Aub, H3K27me3 and H3K9me3 and an active marker H3K4me3 in embryos and ESCs (**Fig. 3l**). Since this model is derived from bulk CUT&RUN analyses, we acknowledge that these data demonstrate genome-wide co-enrichment at the population level and do not directly resolve whether all these histone modifications coexist within the same cells or on the same alleles. Nevertheless, this model emphasizes not only the co-occurrence but also the functional hierarchy of these histone modifications. This model positions H2Aub as a primary repressor, while H3K27me3 and H3K9me3 may play a supportive role and server as reinforcer, explaining how multiple histone markers cooperate to establish and maintain a transcriptionally poised state (**Fig. 3l**).

### H2Aub helps deposit H3K27me3 but restricts H3K4me3 to maintain a polyvalent state

We next sought to investigate the molecular basis underlying polyvalent maintenance. Although PRC1 and PRC2 are known to recruit each other in ESCs ^45, 46^, they play distinct and largely non-redundant role in preimplantation embryos ^39^. However, how these complexes interact to drive polyvalent establishment and maintenance during post-implantation development remains unknown. To address this question, we performed *in vivo* RING1A/1B degradation by dTAG^V^-1 injection at E5.5 (**Fig. 3a, b**) and examining H3K27me3 level at E6.5. Immunostaining showed that H3K27me3 signals were largely lost following H2Aub depletion, suggesting that H2Aub is necessary for H3K27me3 deposition in post-implantation embryos (**Fig. 4a**). Consistent with these observations, CUT&RUN analysis of E6.5 EPI revealed significant reduction of H3K27me3 signals on polyvalent genes (C1 group), such as *Tbx3*, as well as PcG-silenced genes (C3 group) such as *Tbx1* (**Fig. 4b, c**). Interestingly, loss of H2Aub increases H3K4me3 levels at promoters of polyvalent genes (e.g., *Tbx3*), but not at promoters of PcG-silenced genes (e.g., *Tbx1*) (**Fig. 4b, c**). This difference explains why PRC1 loss activates more polyvalent genes (C1) than PcG-silenced genes (C3) (**Fig. 3k**). These findings demonstrate that H2Aub plays a key role in depositing H3K27me3 and restricting H3K4me3 level to maintain the polyvalent chromatin state.

**Figure 4.**
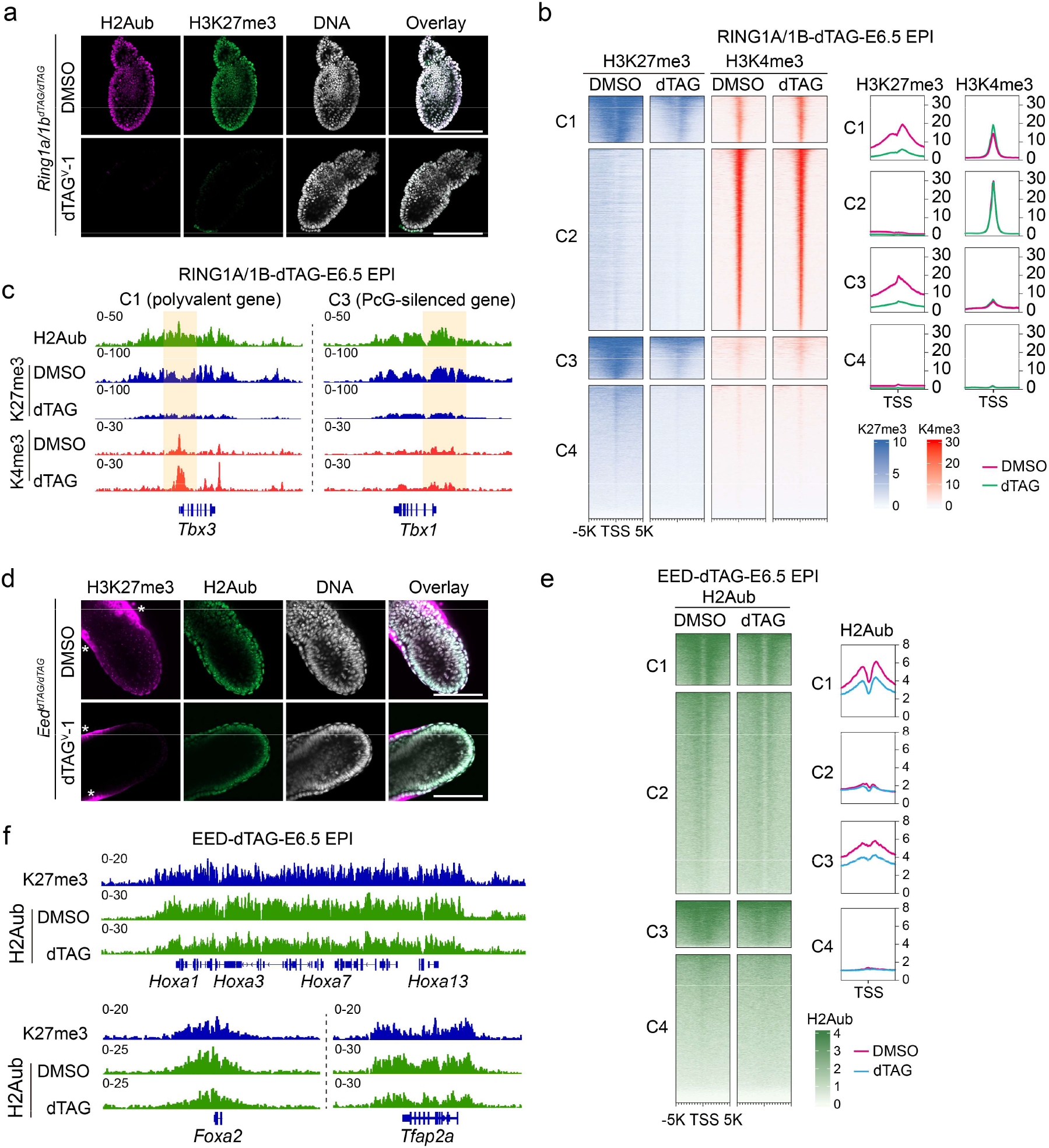
H2Aub is required for H3K27me3 deposition and it restricts H3K4me3 level to maintain polyvalent state. (**a**) Immunostaining showing loss of H3K27me3 following acute depletion of RING1A/RING1B and H2Aub in E6.5 embryos. DMSO or dTAGV-1 was injected into pregnant females at E5.5, and embryos were collected at E6.5. The VE lineage was removed prior to staining to prevent non-specific H3K27me3 signal. DNA was stained with Hoechst 33342. Scale bar, 100 μm. (**b**) Heatmap showing changes in H3K27me3 and H3K4me3 levels following RING1A/1B depletion in E6.5 EPI. The H3K27me3 levels were normalized with E coli. DNA spike-in. (**c**) Genome browser views of H3K27me3 and H3K4me3 levels around *Tbx3* and *Tbx1* genes with or without RING1A/1B depletion in E6.5 EPI. (**d**) Immunostaining of H2Aub following acute depletion of EED and H3K27me3 in E6.5 embryos. DNA was stained with Hoechst 33342. Scale bar, 100 μm. * showed the non-specific staining. (**e**) Heatmap showing the H2Aub changes after acute EED and H3K27me3 depletion in E6.5 EPI. (**f**) Genome browser views of H2Aub levels around *Foxa2* and *Tfap2a* genes with or without EED depletion at E6.5 EPI.

We next examined whether H3K27me3 contributes to H2Aub deposition. To this end, we performed *in vivo* EED degradation by dTAG^V^-1 injection at E5.5 (**Fig. 1c**) and analyzed H2Aub level at E6.5. Immunostaining showed that although H3K27me3 was completely depleted, H2Aub levels were only slightly reduced (**Fig. 4d**). CUT&RUN analysis further confirmed this result, showing that H2Aub signals were largely maintained at polyvalent genes (C1 group, e.g., *Hoxa1*, *Hoxa3*, *Foxa2*, and *Tfap2a*) at E6.5 (**Fig. 4e, f**). These results suggest that H3K27me3 only plays a minor role in H2Aub deposition on polyvalent genes. In summary, our data suggest that H2Aub deposition is upstream of H3K27me3 in the establishment of polyvalent chromatin state during post-implantation development.

### Polyvalency establishment and dynamics during pre- to post-implantation development

Bivalency was previously thought to be established in post-implantation embryos ^47^. However, our recent study ^40^, together with others ^48^, suggests that the bivalency has already started to be established in peri-implantation embryos. To investigate how the polyvalency is established during embryonic development, we preformed H3K4me3, H2Aub, H3K27me3 and H3K9me3 profiling across multiple developmental stages and lineages, including E4.5 inner cell mass (ICM), E7.5 mesoderm (Mes), ectoderm (Ect) and endoderm (End) (**Fig. 5a and Extended Data Fig. 5a**). Hierarchical clustering and principal component analysis (PCA) revealed that different histone markers exhibit different relationship among the different cell states (**Fig. 5b, c**). In addition, compared to H3K27me3, H2Aub and H3K9me3 showed smaller variation across the cell lineages (**Fig. 5c**), suggesting H2Aub and H3K9me3 chromatin landscape showed lower global variation across lineages. Similar to E6.5 cells where most bivalent genes are polyvalent (**Fig. 2c, d**), the majority of bivalent genes are also polyvalent in E4.5 and E7.5 cells (**Extended Data Fig. 5b, c**). Although the total number of polyvalent genes exhibit an increased trend from E4.5 ICM to E6.5 EPI and E7.5 Mes and End (**Fig. 5d**), the gene numbers are variable (**Fig. 5e**), suggesting a progressive polyvalency establishment, maintenance and resolution across the developmental stages.

**Figure 5.**
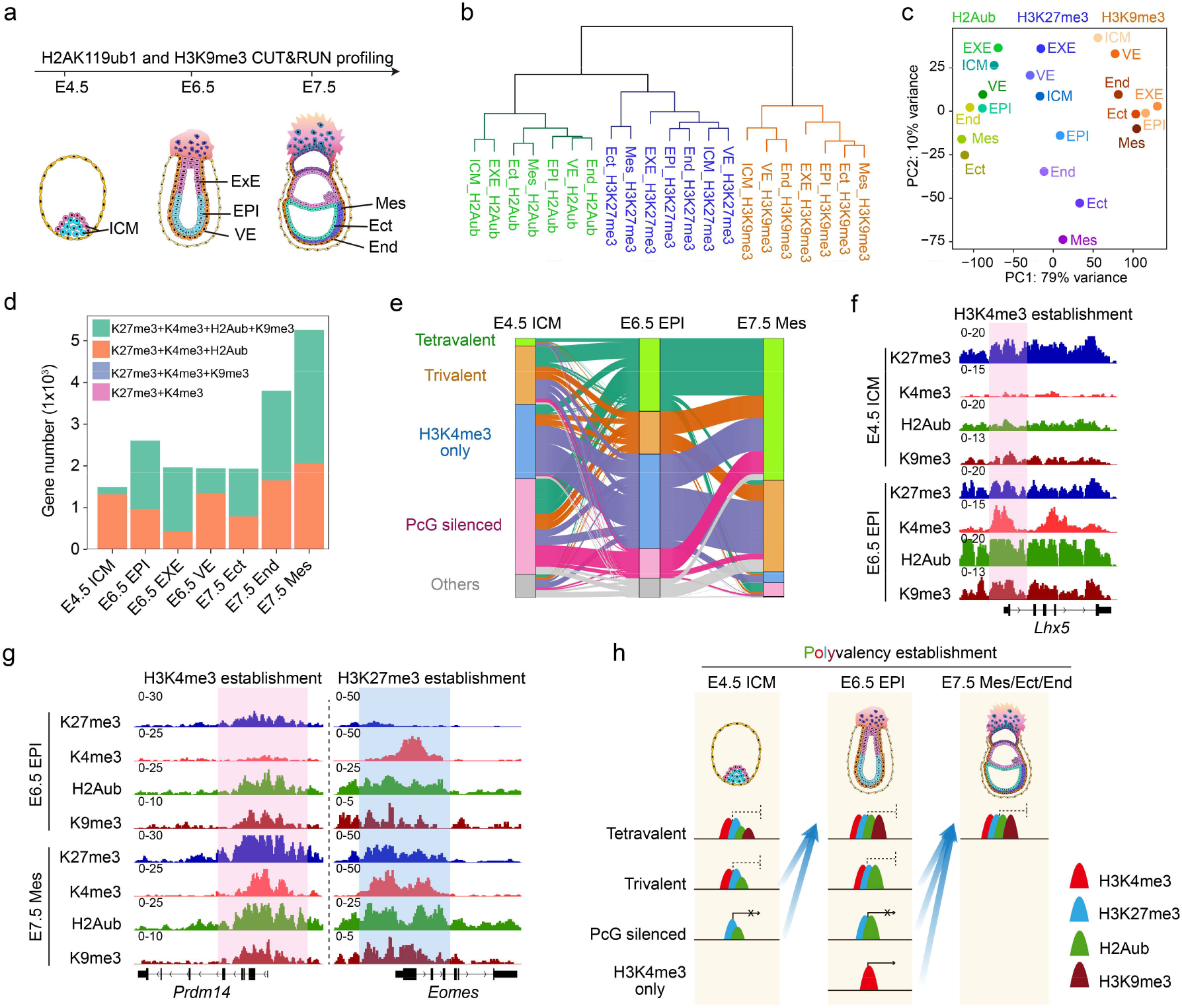
Dynamics of polyvalency from pre- to post-implantation development. (**a**) Schematic illustrating H2Aub and H3K9me3 CUT&RUN profiling across developmental stages. ICM, inner cell mass; ExE, extraembryonic ectoderm; EPI, epiblast; VE, visceral endoderm; Mes, mesoderm; Ect, ectoderm; End, endoderm. (**b**) Hierarchical clustering of H2Aub, H3K27me3, and H3K9me3 profiles across developmental stages. (**c**) Principal component analysis (PCA) of H2Aub, H3K27me3, and H3K9me3 profiles across developmental stages. (**d**) Bar plot showing the number of genes marked by different combinations of H3K27me3, H3K4me3, H2Aub, and H3K9me3 across embryonic lineages at the indicated stages. (**e**) Sankey diagram showing transitions in gene chromatin-state categories between E4.5 ICM, E6.5 EPI and E7.5 Mes. Genes are classified as tetravalent, trivalent, H3K4me3-only, PcG-silenced, or unmarked. The widths indicate the number of genes transitioning between categories. Only transitions involving tetravalent/trivalent in at least one stage were plotted. (**f**) Genome browser views illustrating establishment of polyvalency from E4.5 ICM to E6.5 EPI through acquisition of H3K4me3. (**g**) Genome browser views illustrating establishment of polyvalency from E6.5 EPI to E7.5 Mes through acquisition of H3K4me3 or H3K27me3. (**h**) Schematic model illustrating dynamic regulation of polyvalency during pre- to post-implantation development.

We next analyzed the dynamics of polyvalency during the developmental process. Surprisingly, only 11.41% of polyvalent genes are at the tetravalent state (K27me3/K4me3/H2Aub/K9me3) in E4.5 ICM (**Fig. 5d and Extended Data Fig. 5c)**, which increases to 62.97% at E6.5 EPI (**Fig. 5d and Fig. 2c**), suggesting a quick shift from a trivalent (K27me3/K4me3/H2Aub) to a tetravalent state (**Extended Data Fig. 6a**), likely due to re-establishment of H3K9me3 at promoters in post-implantation embryos ^49^. In addition, many tetravalent genes (48.75%, 800 of 1,641) are established de novo from the PcG-silenced state (**Fig. 5e, Extended Data Fig. 6a**). For example, *Lhx5* is completely silenced by PcG at E4.5 ICM, but acquiring polyvalent state by gaining H3K4me3 at E6.5 EPI (**Fig. 5f**). On the other hand, polyvalency can be resolved during E4.5 ICM to E6.5 EPI transition through the removal of the H3K27me3 mark (**Fig. 5e, Extended Data Fig. 6a**). Unlike the E4.5 ICM to E6.5 EPI transition, polyvalency resolution from E6.5 EPI to E7.5 Mes/Ect/End is mainly achieved by the removal of H3K4me3 (**Extended Data Fig. 6b-d,** resolved), while the gained polyvalency is due to acquisition of either H3K4me3 or H3k27me3 (**Fig. 5e, g and Extended Data Fig. 6e, f**).

Collectively, these developmental stage-specific analyses provide a comprehensive view of how polyvalency is established during peri- to post-implantation embryos development. Building on our previous bivalency model at E4.5 ICM ^40^, we now show that the chromatin state previously described as bivalent is largely trivalent, indicating that trivalency is directly established in the E4.5 ICM (**Fig. 5h**). This trivalent state subsequently acquires H3K9me3 to form tetravalency at the E6.5 EPI (**Fig. 5h**). As development proceeds, the polyvalency undergoes dynamic resolution, maintenance, or establishment that collectively shaping the chromatin landscape during gastrulation.

### PRC1 plays a key role in organogenesis but not in gastrulation

Although bivalency has been extensively described and characterized at the epigenomic level ^21, 50^, much less is known about how bivalency (redefined as polyvalency) functionally contributes to biological processes, particularly *in vivo* at the organism level. We next sought to investigate whether the PRC1-mediated polyvalency contributes to embryonic development. Consistent with previous observations that bivalent genes are predominantly developmental regulators ^21^, the majority of polyvalent genes are associated with developmental processes (**Extended Data Fig. 7a**). Gene ontology (GO) analysis further suggest that the polyvalent genes are enriched in pathways related to organogenesis but much less for gastrulation (**Extended Data Fig. 7a**), suggesting the potential roles of polyvalency in regulating organogenesis.

Maternal *Ring1a* and *Ring1b* double knockout or zygotic H2Aub depletion led to 2-cell or 4-cell arrest ^38, 39^, making it impossible to study PRC1 function in post-implantation development. Taking advantage of our dTAG mice, we were able to rapidly deplete PRC1 and H2Aub in embryos at defined developmental stages *in vivo* (**Fig. 3b**). By continuously administering dTAG^V^-1 beginning at E6.0, we depleted RING1A/1B and collected embryos at E7.5, E8.5 and E9.5 for morphological analysis (dTAG#1) (**Fig. 6a and Extended Data Fig. 7b**). Strikingly, RING1A/1B depletion did not cause obvious morphological abnormalities at E7.5 or E8.5. However, E9.5 embryos were markedly smaller and exhibited a typical failure of embryonic turning, a phenotype strongly associated with defects in somite formation, paraxial mesoderm development or neural tube closure (**Fig. 6b**) ^51, 52^. Given that gastrulation is a prerequisite for organogenesis ^53^, we next attempt to define the developmental window when PRC1 exerts its function, while minimizing potential effects on gastrulation. To this end, we initiated continuous dTAG administration starting at E7.0 (dTAG#2), allowing PRC1 and H2Aub to be fully depleted by E7.5, a stage when gastrulation is largely completed (**Fig. 6a**). Interestingly, dTAG#2 treatment resulted in similar morphological defects at E9.5 as that of dTAG#1 (**Fig. 6b**), indicating that the abnormalities are caused by PRC1 loss during early organogenesis.

**Figure 6.**
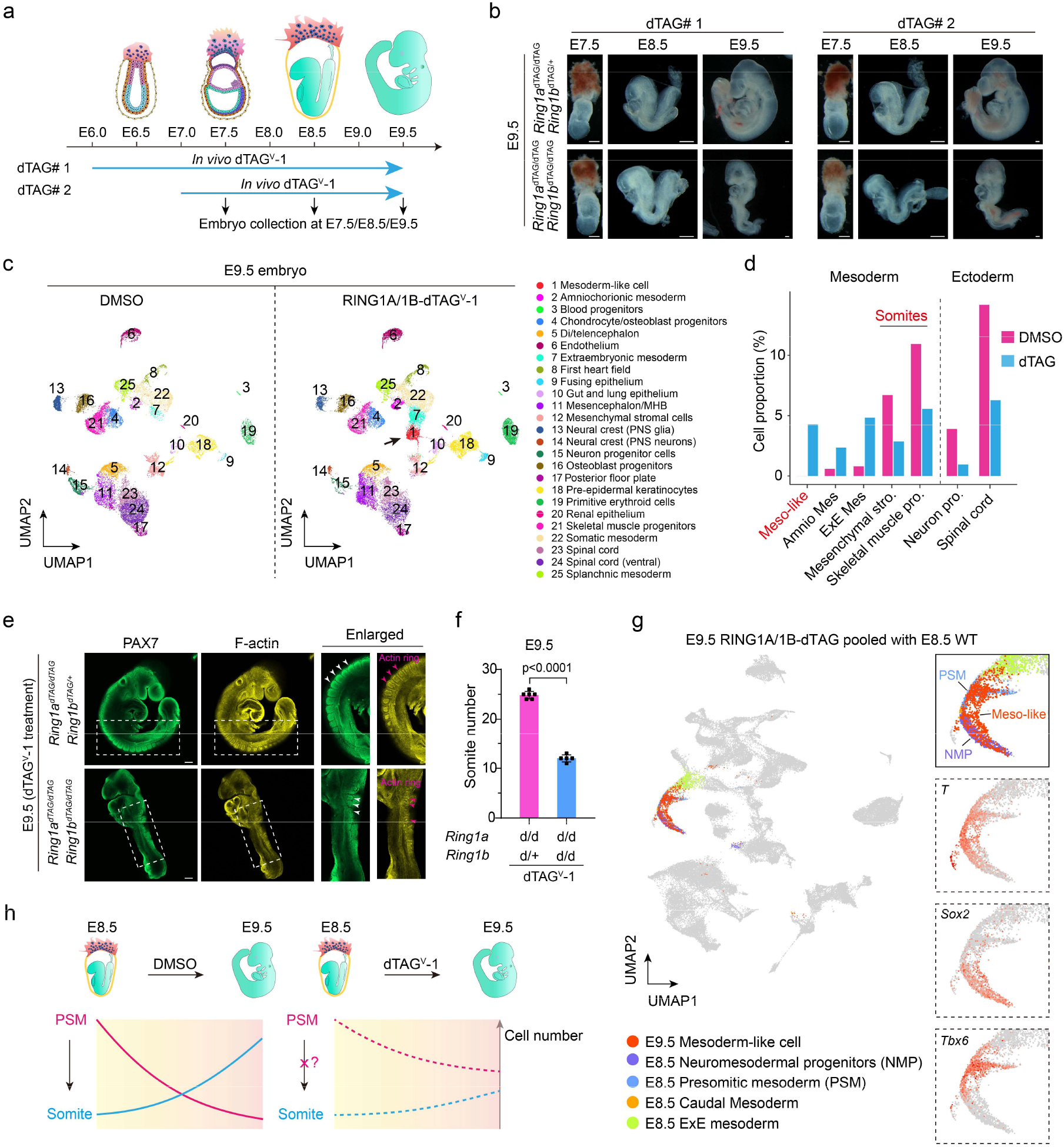
PRC1 degradation results in organogenesis failure. (**a**) Schematic illustration of dTAG^V^-1-mediated *in vivo* RING1A/1B depletion during gastrulation and organogenesis. dTAG^V^-1 was administered by IP injection every 12 h from E6.0 to E9.5 (dTAG#1) or from E7.0 to E9.5 (dTAG#2). Embryos were collected at E7.5, E8.5, and E9.5 for further analysis. (**b**) Images of representative embryos at E7.5, E8.5, and E9.5 with or without RING1A/1B depletion. Within each developmental stage and treatment condition, embryos were derived from the same litter. The RING1A-depleted embryos are used as controls. Images are representative of at least three litters for each developmental stage. Scale bar, 200 μm. (**c**) UMAP visualization of single-cell transcriptomic data from E9.5 embryos with or without RING1A/1B depletion. Each dot represents a single cell, colored by cluster identity. (**d**) Cell proportions corresponding to mesoderm and ectoderm related lineages that showed significant changes after RING1A/1B depletion. (**e**) Immunostaining of E9.5 embryos showing decreased somite number after RING1A/1B depletion. *Ring1a*^dTAG/dTAG^; *Ring1b*^dTAG/+^ and *Ring1a*^dTAG/dTAG^; *Ring1b*^dTAG/dTAG^ embryos are derived from the same litter. The RING1A-depleted embryos are used as controls. PAX7 staining represent the somite (white arrows), and F-actin represent the actin-ring (pink arrows). One representative image from three independent experiments was shown. Scale bar, 400 μm. (**f**) Quantification of somite number in E9.5 embryos following RING1A/1B depletion (n=5). RING1A-depleted embryos (n=6) were used as controls. Embryos analyzed were derived from three independent litters. p values were calculated with Student’s t-test. (**g**) UMAP visualization of integrated single-cell data from E9.5 RING1A/1B-dTAG-treated embryos pooled with E8.5 wild-type (WT) embryos. E9.5 mesoderm-like cells and E8.5 mesoderm-related populations, including neuromesodermal progenitors (NMP), presomitic mesoderm (PSM), caudal mesoderm, and extraembryonic mesoderm (ExE mesoderm) are highlighted. Expression patterns of *T*, *Sox2*, and *Tbx6* are shown on the right. (**h**) Schematic illustration of the developmental trajectories from E8.5 to E9.5 for PSM to somite differentiation, under DMSO or RING1A/1B-dTAG treatment.

### PRC1 depletion results in PSM and NMP arrest

To further determine which cell types are affected by PRC1 loss, we performed single-cell RNA-seq of E9.5 embryos with or without PRC1 degradation (**Fig. 6c**). Consistent with the observed embryonic turning defects, PRC1 depletion resulted in a marked reduction in somite-associated lineages (e.g., mesenchymal stromal cells and skeletal muscle progenitor) and neurogenic lineages (e.g., neuron progenitor and spinal cord), accompanied by an increase in mesoderm-related populations, whereas most other cell types were only minimally affected (**Fig. 6c, d and Extended Data Fig. 7c**). Immunostaining further confirmed that PRC1 loss led to a reduced number of somites (**Fig. 6e, f**). Together, these results indicate that PRC1 is essential for proper somitogenesis.

We next sought to determine the cause of the somitogenesis defects. Notably, dTAG treatment resulted in the appearance of a new cell population that was absent in the DMSO control group (Cluster 1, black arrow, **Fig. 6c**). Marker genes of this cluster of cells suggested they were mesoderm-like cells with *T* expression (**Extended Data Fig. 7d**). Given that somites are generated by the presomitic mesoderm (PSM), and PSM itself is derived from neuromesodermal progenitors (NMP), which are bipotent cells capable of generating either mesodermal or neurogenic lineages during organogenesis (E8.5 and E9.5) ^54–57^, we hypothesize that the mesoderm-like cells represent PSM or NMP cells that fail to progress to somite and instead are arrested from E8.5 to E9.5 following PRC1 depletion.

To test this possibility, we performed an integrated analysis of E8.5 wild-type and E9.5 RING1A/1B-dTAG single-cell RNA-seq datasets. We found that indeed the E9.5 mesoderm-like cells from RING1A/1B-dTAG embryos co-clustered with E8.5 NMP (*T*^+^/*Sox2*^+^) and PSM (*T*^+^/*Tbx6*^+^) populations (**Fig. 6g and Extended Data Fig. 7d**), confirming that these cells represent arrested NMP and PSM. Together, these results indicate that PRC1 depletion blocks PSM and NMP cells from developing into the respective somite lineages, leading to reduced formation of these cell types and the accumulation of PSM and NMP populations (**Fig. 6h**).

### PRC1-driven somitogenesis is independent of *Hox* gene regulation

We next sought to understand the underlying molecular mechanism by which PRC1 regulates PSM development and somitogenesis. To minimize indirect effects caused by prolonged dTAG treatment (PRC1 depletion), we performed single-cell RNA-seq at E8.5 (**Fig. 7a**), a stage when overall embryonic morphology appeared normal (**Fig. 6b**). Single-cell transcriptomic analysis revealed that the majority of cell types, including somite-related lineages (e.g., paraxial mesoderm, pharyngeal mesoderm and cranial mesoderm) were largely unaffected following PRC1 depletion (**Extended Data Fig. 8a**), although the polyvalent genes showed upregulation in all detectable lineages (**Extended Data Fig. 8b**). Consistently, the somite number of E8.5 embryo did not decrease upon PRC1 loss (**Fig. 7b, c**). Notably, NMP and PSM cells exhibited a modest reduction, suggesting a potential role for PRC1 in maintaining these progenitor populations. However, this reduction is unlikely responsible for the failure of somitogenesis as NMP and PSM cells were arrested but did not disappear at E9.5 following PRC1 depletion (**Fig. 6c, d, g and h**).

**Figure 7.**
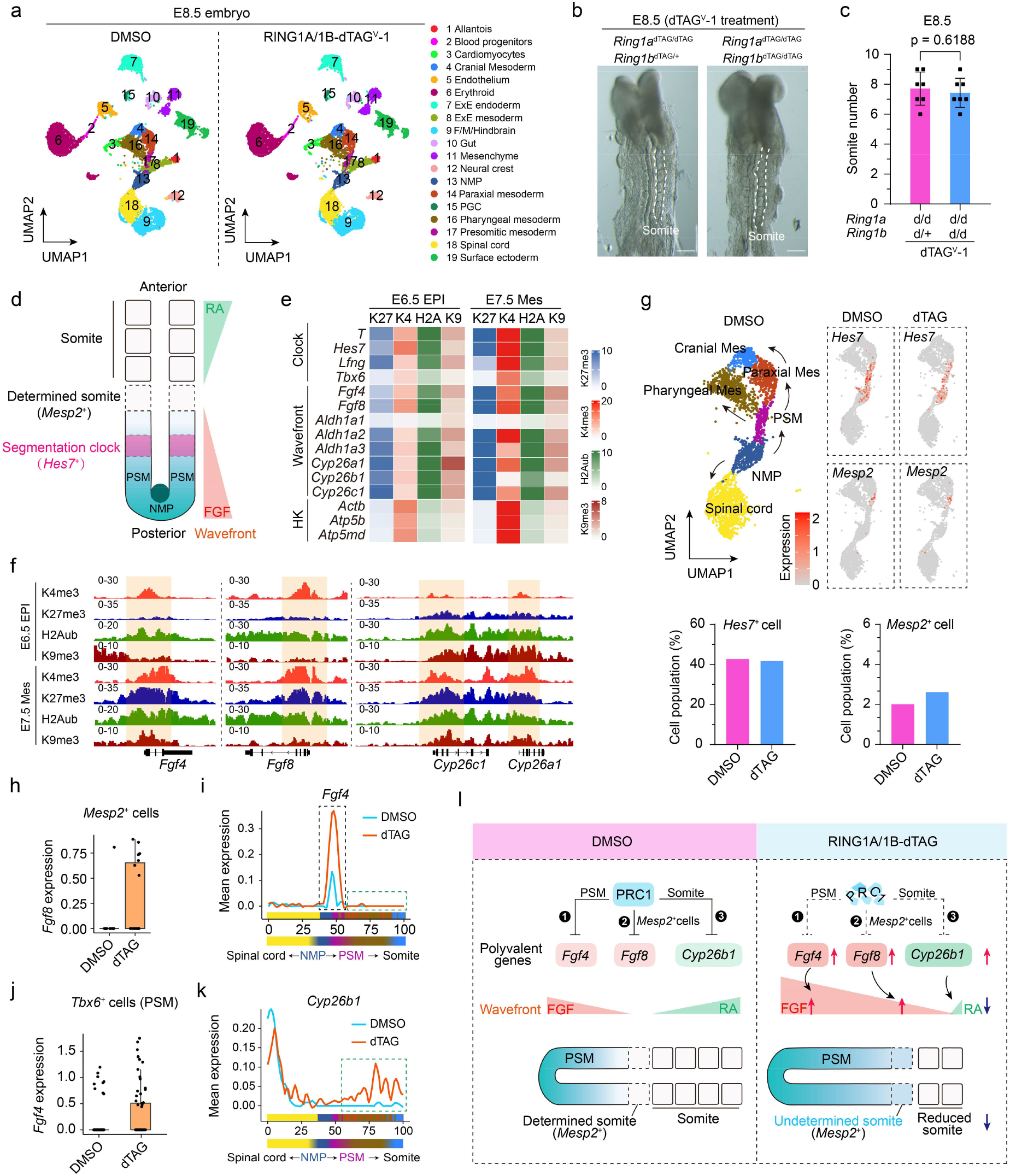
PRC1 regulate organogenesis and polyvalent gene-regulated RA/FGF signaling. (**a**) UMAP visualization of single-cell transcriptomic data from E8.5 embryos with or without RING1A/1B depletion. Each dot represents a single cell, colored by cluster identity. (**b**) RING1A/1B depletion did not affect somite formation at E8.5. *Ring1a*^dTAG/dTAG^; *Ring1b*^dTAG/+^ and *Ring1a*^dTAG/dTAG^; *Ring1b*^dTAG/dTAG^ were obtained from the same litter. The RING1A-depleted embryos were used as controls. Dashed lines indicate somites. One representative image from three independent experiments is shown. Scale bar, 200 μm. (**c**) Quantification of somite number in E8.5 embryos following RING1A/1B depletion (n=7). RING1A-depleted embryos (n=7) are used as controls. Embryos shown are derived from three independent litters. p values were calculated with Student’s t-test. (**d**) Schematic illustration of somite formation along the anteroposterior axis, showing the presomitic mesoderm (PSM), determined somite domain (*Mesp2*⁺), segmentation clock region (*Hes*7⁺), and neuromesodermal progenitors (NMP). Gradients of retinoic acid (RA) and fibroblast growth factor (FGF) are indicated. (**e**) Heatmap showing levels of H3K27me3, H3K4me3, H2Aub and H3K9me3 at representative segmentation clock and wavefront associated genes in E6.5 EPI and E7.5 Mesoderm lineages. Color scales indicate average levels around the TSS for each modification. (**f**) Genome browser views showing representative wavefront related genes in E6.5 EPI and E7.5 Mes cells. (**g**) UMAP visualization of mesoderm related cell populations under DMSO or dTAG treatment. Left up panel, UMAP showing annotated mesodermal subpopulations. Right up panel, feature plots showing *Hes7* and *Mesp2* expression under DMSO or dTAG treatment. Left bottom panel, proportion of *Hes7*⁺ cells within the NMP and PSM. Right down, proportion of *Mesp2*⁺ cells within the paraxial mesoderm. (**h**) RING1A/1B depletion leads to *Fgf8* upregulation in *Mesp2*^+^ cells. (**i**) Mean *Fgf4* expression along a one-dimensional transcriptional ordering delineates the spinal cord-NMP-PSM-somite axis with or without RING1A/1B depletion. Color code as in (g). The green dashed box indicates cell lineages in which *Fgf4* expression is not affected, whereas the green dashed box indicates cell lineages in which *Fgf4* expression is upregulated following RING1A/1B depletion. (**j**) RING1A/1B depletion leads to *Fgf4* upregulation in *Tbx6*^+^ cells. (**k**) Mean expression of *Cyp26b1* along a one-dimensional transcriptional ordering delineates the spinal cord-NMP-PSM-somite axis with or without RING1A/1B depletion. Color code as in (g). The green dashed box indicates cell lineages in which *Cyp26b1* expression is upregulated following RING1A/1B depletion. (**l**) Schematic illustration of how PRC1 may regulate somitogenesis through three mechanisms: (1) Silencing *Fgf4* in PSM. (2) Silencing *Fgf8* in *Mesp2*^+^ cells. (3) Silencing *Cyp26b1* in formed somites.

Given the well-established role of PRC1 in repressing *Hox* gene expression ^58, 59^, and the essential function of *Hox* genes in specifying anteroposterior identity of somites ^60, 61^, we next examined whether PRC1 contributes to somitogenesis, potentially through regulating HOX gene expression. To our surprise, the majority of the *Hox* gene clusters (e.g., *Hoxa, Hoxb, Hoxc* and *Hoxd*) were already expressed in NMP and PSM cells (**Extended Data Fig. 8c, d**), which is consistent with previous study ^62^, suggesting PRC1 does not have a role in silencing *Hox* genes in NMP or PSM cells. Further analysis revealed that PRC1 depletion led to upregulation of multiple *Hox* genes in some lineages such as allantois (*Hoxa3*, *Hoxa5*, *Hoxa9*, *Hoxa11*, *Hoxb9*, *Hoxc8*, *Hoxc10*, *Hoxd3* and *Hoxd8 etc.*) and gut (*Hoxa1*, *Hoxa5*, *Hoxb2* and *Hoxd1 etc.*) (**Extended Data Fig. 8e**). In contrast, most of the *Hox* genes in PSM and NMP were not upregulated following PRC1 depletion, with some even decreased (e.g., *Hoxa3*, *Hoxa7*, *Hoxb4* and *Hoxc4*) (**Extended Data Fig. 8e**), which were most likely due to indirect effects. These results indicate that PRC1 only plays a minor role in silencing *Hox* genes in PSM and NMP cells, and therefore PRC1 regulates early somite formation is unlikely through a *Hox* gene*-*dependent mechanism. This conclusion is consistent with previous results demonstrating that *Hox* genes function primarily in specifying the anteroposterior identity and somite patterning, rather than in controlling somite formation ^60^. Together, these results indicate PRC1-driven early somite formation is independent of *Hox* gene regulation.

### PRC1 regulates FGF signaling and the wavefront through modulating polyvalent genes

Somitogenesis is governed by coordinated action of a molecular segmentation clock and a moving determination front (wavefront) ^57, 63^. The segmentation clock arises from oscillatory transcription driven mainly by Notch and Wnt signaling pathways in PSM. This clock is characterized by the cyclic expression of genes such as *Hes7* and *Lfng*, generating a periodic temporal signal that determines when a new somite should form (**Fig. 7d**) ^57, 63^. The wavefront is established by opposing molecular gradients of high posterior FGF signaling and high anterior retinoic acid (RA) signaling along the PSM. These gradients define a threshold position where FGF activity drops below a critical level, allowing the oscillatory clock to arrest and permitting *Mesp2* activation. This way, temporal information from the segmentation clock is translated into spatially defined somite boundaries (**Fig. 7d**) ^57, 63^.

Notably, the polyvalent chromatin states were detected at both segmentation clock genes (e.g., *T*, *Hes7* and *Lfng*) and wavefront related genes (*Fgf4*, *Fgf8*, *Aldha2* and *Cyp26a1*) in the E6.5 EPI and were maintained in E7.5 mesoderm (**Fig. 7e, f**), suggesting PRC1 may regulate somite formation by controlling segmentation clock and wavefront. To determine whether PRC1 is involved in regulating segmentation clock, we examined the expression of the clock associated genes *Hes7* and *Mesp2* in NMP and PSM related cell types. UMAP analysis showed that PRC1 loss did not significantly alter the expression of these genes (**Fig. 7g**). Importantly, the proportion of *Hes7*⁺ cells (DMSO 42.69% *vs* dTAG 41.67%) and *Mesp2*⁺ cells (DMSO 2.00% *vs* dTAG 2.61%) were not significantly altered upon PRC1 depletion (**Fig. 7g**, right panel). These results collectively indicate that PRC1 loss does not disrupt the segmentation clock.

We next investigated whether PRC1 regulates the wavefront. *Fgf8* is a key component of FGF signaling that controls somite boundary positioning through a posterior-to-anterior gradient ^64, 65^. Overexpression of *Fgf8* maintains PSM cells in an undifferentiated caudal state and blocks somite segmentation ^64^. Interestingly, PRC1 loss led to modest expression of *Fgf8* expression in *Mesp2*^+^ cells (corresponding to the presumptive somite boundary), where it is normally not expressed in control embryos (**Fig. 7h**). Although these cells have already been specified by the segmentation clock, ectopic *Fgf8* expression might block segmentation and cause somite defects ^64, 65^. In addition to *Fgf8*, *Fgf4* can also contribute to wavefront activity ^66^, and ectopic FGF4 signaling has been shown to impair somite formation ^64^. *Fgf4* is normally expressed at low levels in the PSM (**Fig. 7i**, black dashed box), indicating that *Fgf8* is the primary regulator of the FGF gradient. Interestingly, lineage-ordered expression analysis revealed that PRC1 depletion significantly increased *Fgf4* expression specifically in PSM (*Tbx6*⁺) cells (**Fig. 7i, j**), but not in other lineages (**Fig. 7i**, green dashed box), suggesting that PRC1 helps maintain a proper FGF gradient by repressing *Fgf4* in the PSM.

RA signaling is another key regulator of the wavefront. RA is synthesized by the ALDH1A2, which is expressed in somite related lineages (e.g., paraxial mesoderm and pharyngeal mesoderm), but not in the PSM (**Extended Data Fig. 8f**). This spatial restriction is sufficient to repress *Fgf8* expression and prevent its anterior expansion. Conversely, RA is degraded by CYP26A1, which is highly expressed in the PSM but absent from somite-related lineages (**Extended Data Fig. 8f**). Strikingly, PRC1 depletion results in the derepression of RA-degrading enzymes *Cyp26b1* in somite related lineages (**Fig. 7k**, green dashed boxes). Increased RA degradation in these regions likely allows *Fgf8* expression to expand from the PSM into somite-forming mesoderm, consistent with the abnormal *Fgf8* expression observed upon PRC1 loss (**Fig. 7h**). Notably, knockout of the RA-synthesizing enzyme *Raldh2* results in embryonic turning defects, a phenotype similar to what we observed following PRC1 depletion (**Fig. 6b**) ^67^.

Together, these findings demonstrate that PRC1 regulates wavefront and somitogenesis. PRC1 achieves precise regulation of FGF signaling and wavefront positioning by regulating polyvalent gene via three mechanisms: (1) Repressing *Fgf4* in PSM cells to prevent excessive FGF signaling. (2) Silencing *Fgf8* in *Mesp2*⁺ cells to ensure proper exit from the undetermined mesoderm state and allow segmentation to proceed. (3) Repressing *Cyp26b1* in somite-related lineages to maintain RA gradient (**Fig. 7l**).

## Discussion

By integrating temporal epigenomic profiling, here we identified a polyvalent chromatin state with gene promoters marked by H3K4me3, H3K27me3, H2Aub and H3K9me3 in post-implantation embryos. Using the protein degradation dTAG mouse models, we were able to rapidly degrade PRC1 (RING1A/1B-dTAG) or PRC2 (EED-dTAG) to address their functions in regulating the polyvalent genes in a developmental stage-specific manner. We show that H2Aub rather than H3K27me3, is the principal determinant of transcriptional repression at poised promoters of polyvalent genes. Our findings highlight an H2Aub-centric, H3K27me3, H3K9me3 and H3K4me3 involved polyvalency model regulating the poised state. Importantly, due to pre-implantation lethality caused by PRC1/H2Aub depletion ^38, 39^, its role in post-implantation development cannot be determined by conventional approaches. Taking advantage of the dTAG approach, we demonstrate that PRC1 plays a key role in regulating organogenesis, but not gastrulation. We further demonstrate that PRC1 regulates somitogenesis by modulating FGF signaling and the wavefront through silencing polyvalent genes. Together, these findings reveal the molecular mechanism by which PRC1 coordinates somitogenesis through RA/FGF signaling.

PRC1 was considered as a gastrulation regulator, given that the zygotic *Ring1b* knockout led to gastrulation arrest ^68, 69^. However, our results showed that acute depletion of RING1A/1B from E6.0 led to organogenesis but not gastrulation defects, which showed much later defects compared to previous studies. One explanation is that the dTAG system enables rapid depletion of RING1A/1B at the E6.0 stage without disrupting pre-implantation development, whereas the gastrulation defect of the zygotic *Ring1b* knockout embryos is likely caused by the cumulative impact of multiple indirect effects. Indeed, consistent with our dTAG results, tamoxifen induced *Ring1a*/*Ring1b* double knockout caused E8.5 but not E7.5 defects ^70^. Notably, RING1A can partially compensate for RING1B deficiency by preserving residual H2Aub deposition ^6, 38, 71^. Accordingly, *Ring1b* single knockout leads to a partial reduction of H2Aub levels, whereas combined depletion of RING1A and RING1B results in a near-complete loss of H2Aub ^6^. These differences in H2Aub abundance likely result in different transcriptional changes. However, how the different transcriptional alterations contribute to the differential phenotypes remain to be investigated.

The Polycomb silencing system was initially discovered in genetic screens in Drosophila for genes important for body patterning ^72, 73^. The preimplantation lethal phenotype of PRC1 core subunit depletion in mice prevented the study of PRC1 function in gastrulation and organogenesis. By applying the dTAG system, we were able to overcome this technical limitation and uncover a previously inaccessible role of PRC1 in post-implantation development, particularly in somitogenesis. Importantly, we reveal that PRC1 participates in these processes through regulating the RA/FGF signaling pathway by silencing polyvalent genes. This finding establishes a direct link between Polycomb-mediated chromatin repression and the control of key developmental signaling pathways. Importantly, although the wavefront model of somitogenesis is well established, its regulation has been largely attributed to transcriptional feedback within signaling pathways themselves, with little insights into the contribution of chromatin-based mechanisms ^57, 74^. By showing that PRC1 represses polyvalent genes linked to RA/FGF signaling, our data suggest that PRC1-mediated chromatin regulation acts as an additional layer that stabilizes the RA/FGF signaling boundary during the somitogenesis. Together, these findings provide a new perspective on how PRC1 repression is integrated with developmental signaling networks to regulate complex morphogenetic events.

One of the interesting features of the polyvalent chromatin state is the presence of H3K9me3, a histone modification associated with constitutive heterochromatin and stable gene silencing. H3K9me3 is enriched in heterochromatin in pre-implantation embryos, and is established on the gene promoters after implantation ^49^, concomitant with the establishment of polyvalency. The function of promoter H3K9me3 is not completely understood. Similar to H2Aub and H3K27me3, it may function as a form of repression that maintains genes in a transcriptionally silent yet activation-competent state, as it is known to coexist with H3K4me3 in a bivalent chromatin configuration ^42, 75, 76^. In our polyvalency model, similar to H3K27me3, H3K9me3 may serve as a buffering mechanism to limit the activation of poised genes. On the other hand, it may serve as a platform for the recruitment of DNA methylation machinery for stable and permanent silencing. Since the function of H3K9me3 in regulating polyvalent genes has not been directly tested in embryos, it should be regarded as an associated chromatin mark whose significance in this context yet to be determined.

A central finding of this study is that PRC1, rather than PRC2, functions as the dominant repressor at genes previously classified as bivalent genes. Although RING1A/1B-dTAG affects both cPRC1 and vPRC1, we reason that the repressing effect is mainly through vPRC1, because cPRC1 functions by recognizing H3K27me3 while when we deplete H3K27me3, there are limited number of genes affected in E6.5 EPI and VE. Although both PRC1 and PRC2 display extensive co-occupancy at CpG island promoters ^12, 77^, and PRC1 is required for H3K27me3 establishment in ESCs ^14, 45, 78, 79^, the role of PRC1 in silencing bivalent genes has been less studied. PRC2-mediated H3K27me3 has historically been regarded as the defining repressive mark at bivalent promoters ^21, 23^, leading to the prevailing view that PRC2 is the primary silencing machinery on bivalent genes. Although long-term loss of PRC2 results in widespread gene upregulation ^80^, acute depletion of PRC2 does not produce the same effect ^30^. Importantly, the defining feature of poised genes is their capacity for rapid transcriptional activation in response to regulatory cues. Our study supports that PRC1 is the dominant repressive regulator associated with the poised transcriptional state, whereas PRC2 functions to reinforce and stabilize the repressed chromatin state.

## Methods

### dTAG mice generation

All experiments were conducted in accordance with the National Institute of Health Guide for Care and Use of Laboratory Animals and were approved by the Institutional Animal Care and Use Committee (IACUC) of Boston Children’s Hospital and Harvard Medical School (protocol number IS00000270-9). Mice were maintained under specific pathogen-free conditions in a controlled environment with temperatures maintained at 20-22 °C, relative humidity of 40–70%, and a 12-hour alternating light/dark cycle. *Ring1a* and *Ring1b* dTAG mouse lines were generated separately using CRISPR/Cas9-mediated genome editing. A solution containing *Ring1a* or *Ring1b* donor DNA (30 ng/μl), sgRNA (30 ng/μl), and *Cas9* mRNA (100 ng/μl) was microinjected into embryos at the early two-cell stage using a Piezo-assisted micromanipulation system (Primer Tech, Ibaraki, Japan). Injected embryos were incubated in KSOM medium for 6 hours before surgical transfer into the oviducts of pseudo-pregnant ICR recipient females (Charles River). Founder (F0) animals were subsequently crossed with C57BL/6J wild-type mice for at least two successive generations to ensure stable germline transmission. *Ring1a*^dTAG/+^ mice were crossed with *Ring1b*^dTAG/+^ mice to generate *Ring1a*^dTAG/dTAG^; *Ring1b*^dTAG/+^ mice for further experiments. Primer sequences used for genotyping are provided in **Supplementary Table 4**.

### *In vitro* fertilization and pre-implantation embryo culture

Superovulation was induced in 7-8-week-old wild-type BDF1 female mice by intraperitoneal administration of 7.5 IU pregnant mare serum gonadotropin (PMSG; BioVendor, RP1782725000), followed 48 hours later by an injection of 7.5 IU human chorionic gonadotropin (hCG; Sigma, C1063). Oocyte-cumulus complexes (OCCs) were harvested 14 hours after hCG treatment. Prior to OCC collection, spermatozoa were isolated from the cauda epididymis of sexually mature male mice (8-12 weeks old) and capacitated in human tubal fluid (HTF) medium (Millipore, MR-070-D) for 40 minutes. Retrieved OCCs were then inseminated with capacitated sperm and incubated together in HTF medium for 6 hours. Zygotes displaying two pronuclei were selected and cultured in KSOM medium (Millipore, MR-106-D) for further development.

### Embryo dissection and cell isolation

To isolate inner cell masses (ICMs) from E4.5 embryos, zona pellucida were removed using Acidic Tyrode’s solution (Millipore). Zona-free embryos were subsequently incubated with anti-mouse serum (diluted 1:3 in KSOM medium; Sigma-Aldrich, M5774) at 37 °C for 30 minutes, followed by exposure to guinea pig complement (1:3 dilution in KSOM; Millipore, S1639) for an additional 30 minutes at 37 °C. Trophectoderm cells were selectively lysed, and the remaining ICMs were mechanically collected using a glass capillary. For E6.5 and E7.5 embryo isolation, embryos were manually isolated from the decidual tissue and the Reichert’s membrane was carefully removed. Embryos were then subjected to enzymatic treatment in 100 μl of 0.25% trypsin supplemented with 2.5% pancreatin (Sigma, P3292) for 15 minutes on ice. Digestion was terminated by the addition of an equal volume of medium containing 10% fetal bovine serum (FBS). For E6.5 embryo, the visceral endoderm (VE) layer was manually stripped away using a glass pipette, after which the epiblast (EPI) and extraembryonic ectoderm (ExE) were dissected using a fine needle under a stereomicroscope. For E7.5 embryos, the extraembryonic region was removed, and the endoderm was first dissected using a fine needle, followed by dissection of the mesoderm and ectoderm. For single-cell isolation at E8.5 and E9.5, embryos were dissociated in 500 μl of TrypLE (Thermo Fisher, 12-605-010) for 10 min at 37 °C. During digestion, samples were gently pipetted every 5 min to facilitate complete dissociation. The reaction was terminated by adding 1.5 ml medium containing 10% FBS. The resulting cell suspension was filtered through a 40 μm cell strainer to remove undigested tissue and debris, and the filtered cells were immediately used for single-cell RNA-seq library preparation.

### Protein degradation *in vivo*

To achieve EED depletion in post-implantation embryos *in vivo*, *Eed*^dTAG/+^ female mice were used to mate with *Eed*^dTAG/dTAG^ male mice, and subjected to intraperitoneal (IP) injection of dTAG^V^-1. For RING1A and RING1B degradation, *Ring1a*^dTAG/dTAG^; *Ring1b*^dTAG/+^ female mice were used to mate with *Ring1a*^dTAG/dTAG^; *Ring1b*^dTAG/+^ male mice, and subjected to IP injection of dTAG^V^-1. For preparation of the dosing solution, 1 mg of dTAG^V^-1 was first dissolved in 25 μl dimethyl sulfoxide (DMSO) and then brought to a final volume of 500 μl by dilution with 10% castor oil (Sigma, C5135). Female mice (∼28 g) were injected with 1 mg dTAG^V^-1 at 12-hour intervals to ensure continuous target protein degradation.

### Immunostaining and confocal microscopy

Blastocyst and post-implantation embryos were fixed in 4% paraformaldehyde and 0.5% Triton X-100 for 20 minutes. After fixation, samples were washed three times with PBS supplemented with 0.1% Triton X-100, followed by blocking in PBS containing 1% bovine serum albumin (BSA) and 0.1% Triton X-100 for 1 hour at room temperature. Embryos were then incubated with primary antibodies diluted in blocking buffer overnight at 4 °C. The antibodies and dilutions used were as follows: anti-H2Aub (1:500, CST, 8240S), anti-EED (1:500, CST, 85322S), anti-HA (1:500, CST, 2367S), anti-H3K27me3 (1:200, Active Motif, 61017) and anti-PAX7 (1:500, Proteintech, 20570-1-AP). Following primary antibody incubation, embryos were washed three times and subsequently incubated with secondary antibodies for 1 hour at room temperature. Secondary antibodies included donkey anti-mouse IgG-Alexa Fluor 568 (1:500, Invitrogen, A10037) and donkey anti-rabbit IgG (H+L)-Alexa Fluor 488 (1:500, Invitrogen, A-21206). DNA was stained with Hoechst 33342 (10 μg/ml, Sigma). Fluorescent images were acquired using a Zeiss LSM800 confocal microscope.

### CUT&RUN, poly(A)-RNA-seq and single-cell RNA-seq library preparation and sequencing

CUT&RUN was performed as previously described ^37^. Briefly, isolated embryos were incubated with Concanavalin A-coated magnetic beads (Polysciences, 86057-3) for 10 mins and then exposed to primary antibodies (anti-H3K4me3, 1:100, CST, 9727S; anti-H3K27me3,1:100, Diagenode, C15410069; anti-H2AK119ub1, 1:100, CST, 8240S; anti-H3K9me3, 1:100, Active Motif, 39062) at 4 °C for overnight incubation. Samples were washed three times, and then incubated with protein A-micrococcal nuclease (pA-MNase, 3 ng/μl) for 2 hours at 4 °C. Subsequently, activation of pA-MNase was performed by incubation with 200 μl of pre-chilled 2 mM CaCl₂ for 20 min at 4 °C, and the reaction was quenched by the addition of 23 μl of 10× stop buffer. To release DNA fragments, samples were incubated at 37 °C for 15 min, followed by the addition of 2.5 μl of 10% SDS and 2.5 μl of Proteinase K (20 mg/ml; Thermo Fisher, AM2546), and subsequently incubated at 55 °C for 1 h. Phenol-chloroform extraction was used for DNA purification. Sequencing libraries were constructed using the NEBNext Ultra II DNA Library Preparation Kit (New England Biolabs, E7645S).

For poly(A)-RNA-seq, isolated embryos were collected and snap-frozen at -80 °C. Full-length cDNA was generated using the SMART-Seq v4 Kit (Clontech, 634890), followed by library construction with the Nextera XT DNA Sample Preparation Kit (Illumina, FC-131-1024). Single-cell RNA-seq libraries were prepared using the Chromium Next GEM Single Cell 3ʹ Kit v3.1 (10x Genomics, 1000269), following the manufacturer’s instructions. All libraries were sequenced on the Illumina NextSeq 1000 platform.

### RNA-seq data analysis

RNA-seq data was processed as described previousl*y* ^37^. R package DESeq2 (v1.46.0) ^81^ was used to calculate differential expressed genes (DEGs). We used all the genes for DEGs calculation without filtering genes with low read counts. Genes with at least 2-fold change and adjusted p-value less than 0.05 were considered as DEGs. R package clusterProfiler (v4.14.3) ^82^ was used for the GO enrichment analysis.

### CUT&RUN data analysis

For CUT&RUN sequencing data, the reads were trimmed by Trimmomatic (v0.39) ^83^ to remove sequencing adaptors. Then the reads were mapped to GRCm38 or GRCh38 reference genome using bowtie2 (v2.4.2) ^84^ with parameters: --local --very-sensitive-local --no-unal --no-mixed --no-discordant --dovetail -I 10 -X 700 --soft-clipped-unmapped-tlen. PCR duplicate reads were removed with Picard MarkDuplicates (v2.23.4) and only proper paired reads with a minimum mapping quality of 30 were retained. For H3K4me3, the peaks were called with MACS2 (v2.2.7.1) ^85^ and the reproducible peaks between two replicates were calculated with the irreproducible discovery rate (IDR) framework ^86^, with the IDR cutoff of 0.05. The FPKM (fragments per kilobase region per million mapped fragments) signal tracks were generated with Deeptools ^87^ bamCoverage (v3.5.1) with 100 bp bin size. The heatmaps of CUT&RUN signals around TSS were generated with Deeptools computeMatrix (v3.5.1) with bin size of 10, and plotted with R packages profileplyr (v1.22.0) and EnrichedHeatmap (v1.36.0). To compare the signals of one histone modification among different stages, the FPKM signals were normalized using the csaw package (v1.40.0) ^88^, with the normalization scale factor calculated by normFactors function using reads counts in 5 kb bin as input. For CUT&RUN with E. coli spike-in, the sequencing reads were mapped to E. coli genome the same way described above and the number of mapped reads was counted. Then the raw reads count mapped to mouse genome in each bin were divided by the total number of reads mapped to E. coli genome and used as the spike-in normalized signal values.

### ChromHMM and clustering analysis

ChromHMM (v1.26) ^89^ was used to model the dynamics of H2Aub among developmental stages from E4.5 to E7.5, which split genomic regions into 7 categories based on the H2Aub signals. We first calculated the signals in each 5 kb bin of the mouse genome with BinarizeBam command using all H2Aub aligned reads bam files as input, and requiring the signal-to-background ratio at least 3-fold (option -f 3). The segmentation model was calculated with LearnModel command using the above calculated 5 kb bin signals as input.

To perform the PCA analysis of CUT&RUN signals, the read counts in 5 kb bins were calculated and log transformed using rlog function in DESeq2 package. Then the prcomp function in R was used to calculate the principal components, with top 10,000 variable regions as input. To perform the clustering analysis, the hclust function in R was employed, with the distance calculated based on the PCA results as input.

### Bivalent/polyvalent gene identification

To identify bivalent genes which were marked by both H3K4me3 and H3K27me3 in each stage, the H3K4me3 peaks were first obtained as described above. Then the average FPKM of H3K27me3 at these regions (±2 kb around H3K4me3 peak center) was calculated and regions with a value at least 3 were considered as strong H3K27me3 domains. Genes with promoter H3K4me3 peaks and strong H3K27me3 domains were identified as bivalent genes. To identify polyvalent genes, the average H2Aub / H3K9me3 signals around the TSS (±2.5 kb) were calculated and regions with average signal greater than 1.5 fold of the trimmed means of H2Aub / H3K9me3 were considered as strong H2Aub /H3K9me3 domains. The trimmed means of H2Aub / H3K9me3 were obtained by first ranking the signals in all the 5 kb bins across mouse genome then removing the top 30% and the bottom 30% bins, the average signal in the remaining bins was the trimmed means of H2Aub / H3K9me3.

### scRNA-seq data analysis

The 10X scRNA-seq data were processed by cellranger (v8.0.1) and analyzed with R package Seurat (v5.1.0) ^90^. The read counts were log-normalized and scaled. Then PCA analysis was performed with RunPCA function in Seurat. Multiple samples of different conditions and different replicates were integrated using Harmony (v1.2.1) ^91^ method with IntegrateLayers function in Seurat. The UMAP (Uniform Manifold Approximation and Projection) analysis was performed using RunUMAP function with top 20 dimensions in Harmony integrated results. Cell clustering was performed using FindClusters function with resolution of 1. To annotate the cell types for each cluster, we first performed label transferring by mapping to the reference dataset using FindTransferAnchors and TransferData functions, then manually checked the cell type assignment for each cluster based on the marker genes of the cluster. For E8.5 embryos, the mouse E8.5 scRNA-seq dataset from R package MouseGastrulationData (v1.20.0) was used as the reference dataset ^33^. For E9.5 embryos, the mouse E9.5 scRNA-seq dataset from TOME project ^92^ (https://tome.gs.washington.edu) was used as the reference dataset. The marker genes of each cluster were identified using FindAllMarkers function in Seurat with minimum expression percentage of 0.5. The pseudo-time analysis was performed with learn_graph and order_cells functions in monocle3 (v1.3.7) ^93^ package.

### Public data used in this study

The nascent RNA TT-seq results of H3K9 methyltransferases dTKO were downloaded from GEO GSE233040 and the calculated average length-normalized results were used ^44^. The H3K9me3 ChIP-seq in mESC data ^49^ were from GSE97778. The H3K4me3 ChIP-seq data in E7.5 embryos ^26^ were from GSE125318.

## Acknowledgements

We thank Drs. Qianying Yang, Yota Hagihara, Shan Jiang and Boyan Wang for their valuable comments on the manuscript. This project was supported by NIH (R01HD116750) and the HHMI. Y.Z. is an investigator of the Howard Hughes Medical Institute. Z.C. is supported by the Trustee award from Cincinnati Children’s Hospital Medical Center and NIH (R00HD104902, R01HD118540).

## Author contributions

Y.Z. conceived and supervised the project. C.Z. designed and performed experiments. Z.C. established the RING1B-dTAG mice. M.W. performed all the bioinformatic analysis. C.Z., M.W., Z.C. and Y.Z. wrote the manuscript. All authors interpreted the data and reviewed the manuscript.

## Data and materials availability

All data generated in this study have been deposited to the NCBI Gene Expression Omnibus (GEO) with accession number GSE316005 and GSE298615.

## Extended Data Figures

**Extended Data Figure 1.**
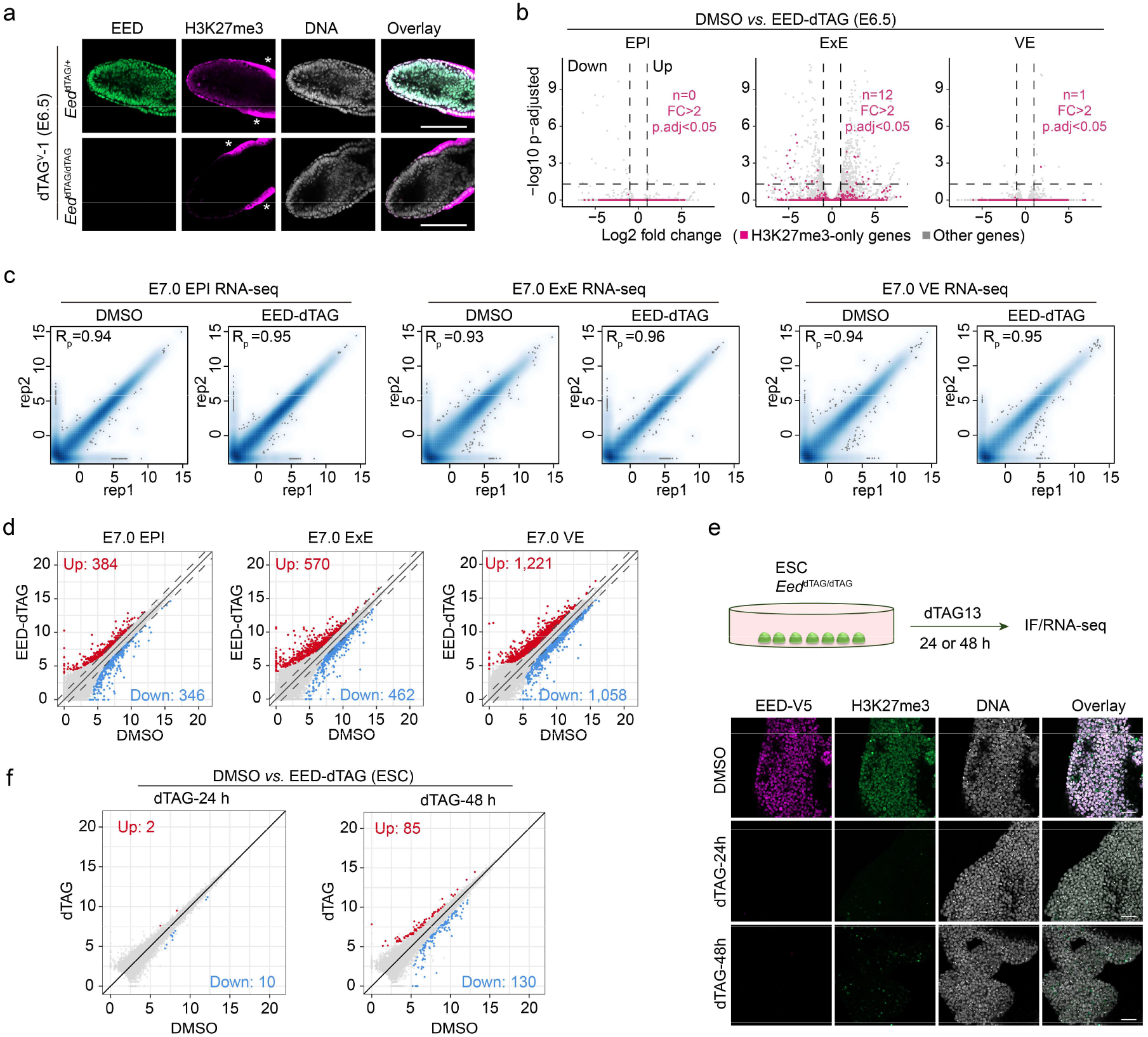
Acute EED depletion results in upregulation of a small group of bivalent genes. (**a**) Immunostaining showing rapid depletion of EED and H3K27me3 in E6.5 embryos *in vivo* following dTAG^V^-1 injection. DNA was stained with Hoechst 33342. Scale bar, 100 μm. Asterisks denote non-specific staining. (**b**) Volcano plots comparing gene expression changes between DMSO and EED-depleted E6.5 EPI, ExE and VE lineages. Red dots represent H3K27me3-only marked genes, and gray dots represent other genes. FC: fold change. (**c**) Scatter plots showing the reproducibility between biological replicates for E7.0 EPI, ExE, and VE RNA-seq datasets. Pearson correlation coefficients are shown. (**d**) Scatter plot comparing gene expression changes between DMSO and dTAG treated E7.0 EPI, ExE and VE. Red and blue dots represent up- and down-regulated repeat subfamilies in the dTAG group, respectively. (**e**) Immunostaining showing EED and H3K27me3 depletion in ESCs with dTAG13 treatment for 24 and 48 h. DNA was stained with Hoechst 33342. Scale bar, 50 μm. (**f**) Scatter plots comparing gene expression changes between DMSO and dTAG groups in ESCs. Red and blue dots represent up- and down-regulated repeat subfamilies in the dTAG group, respectively.

**Extended Data Figure 2.**
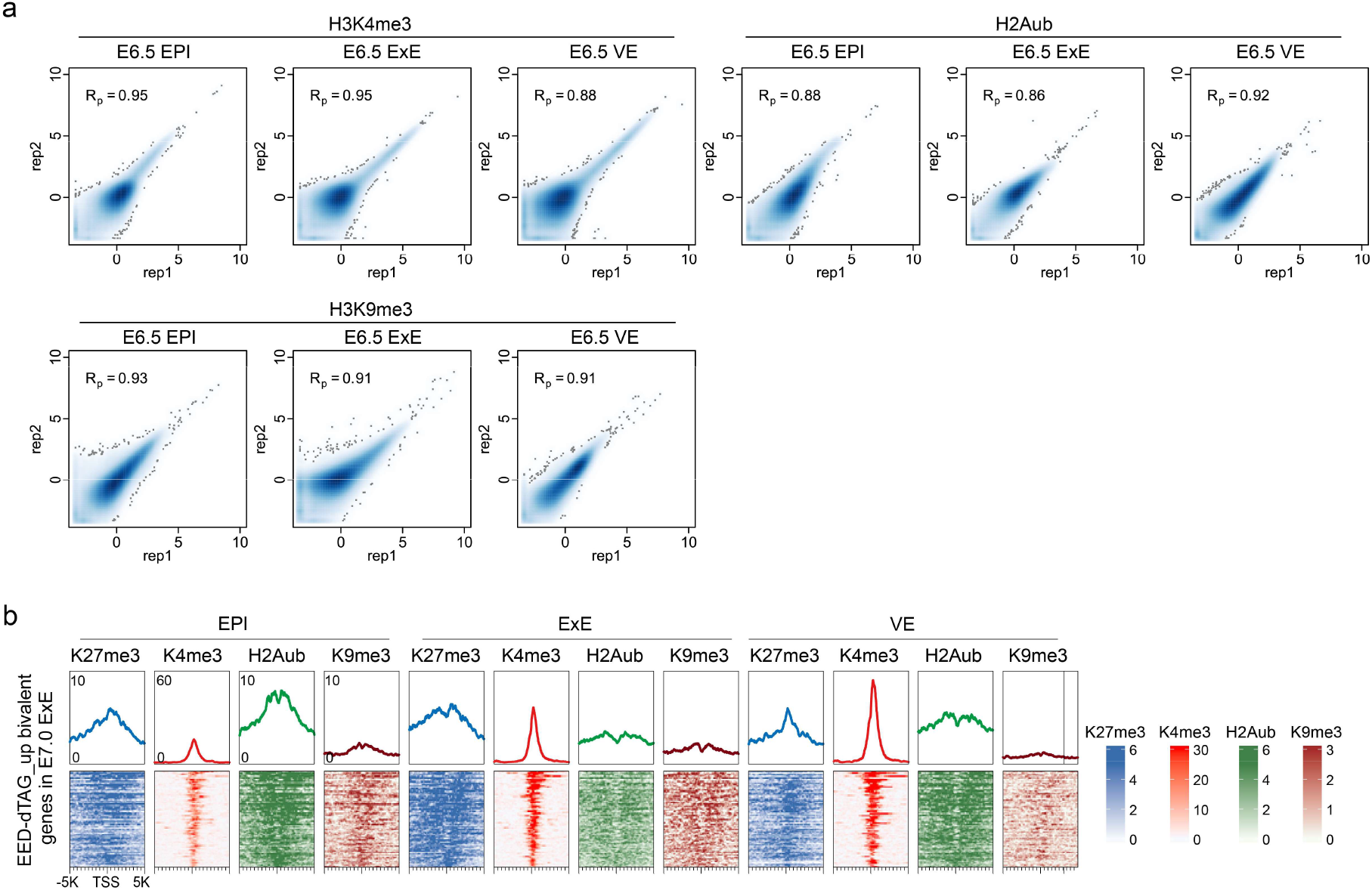
H3K4me3, H2Aub and H3K9me3 profiling in E6.5 embryos. (**a**) Scatter plots showing the reproducibility between biological replicates for H3K4me3, H2Aub and H3K9me3 CUT&RUN datasets. Pearson correlation coefficient is shown. (**b**) Heatmaps and meta plots showing H3K27me3, H3K4me3, H2Aub and H3K9me3 enrichment on bivalent genes that showed upregulation in E7.0 ExE (n=113), but not in E7.0 EPI or VE lineage cells following EED degradation.

**Extended Data Figure 3.**
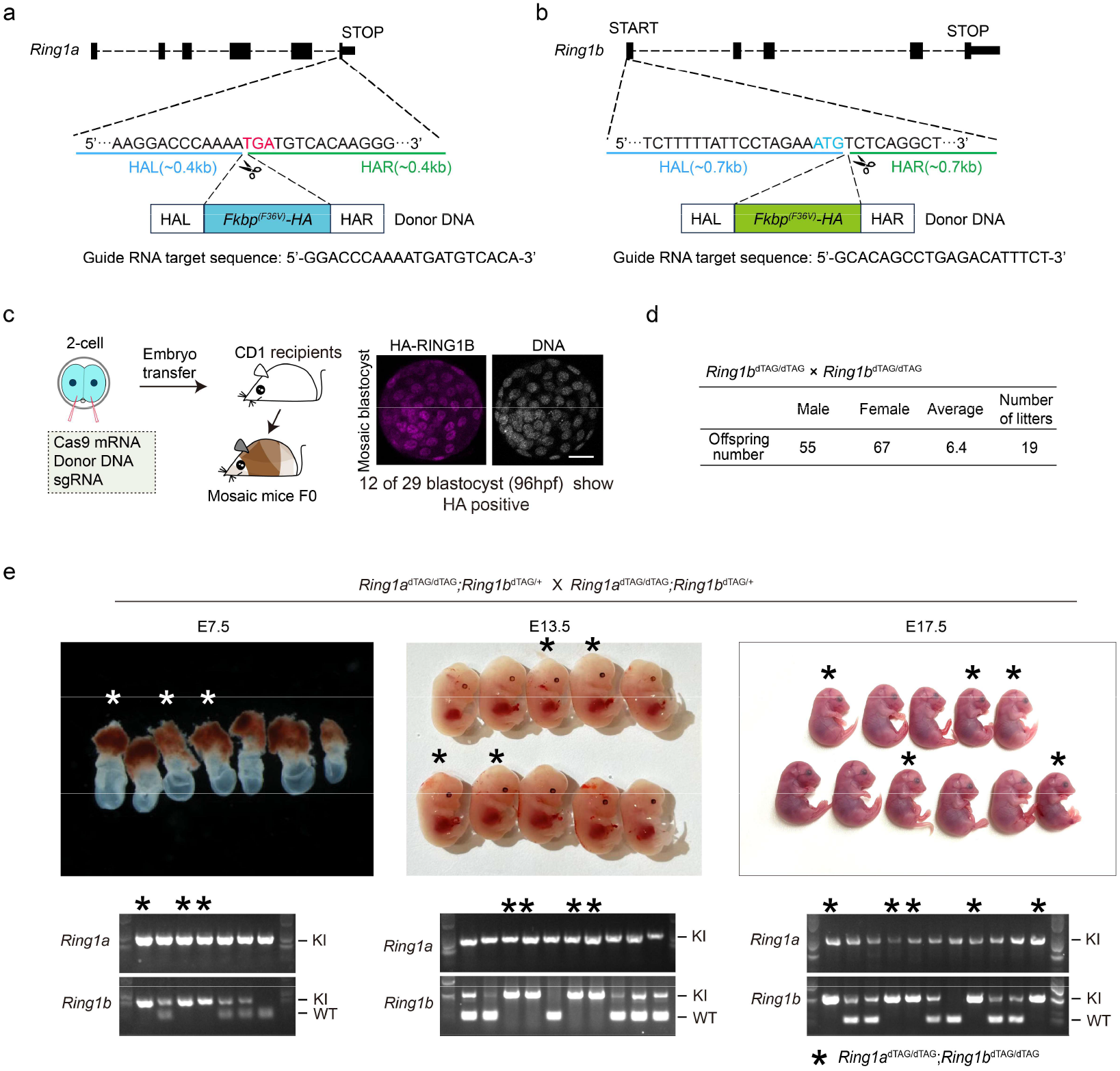
Generation of *Ring1a* and *Ring1b* dTAG mice. (**a, b**) Schematic illustration of the strategy in generating the dTAG knock-in *Ring1a* (a) and *Ring1b* (b) locus. HAL: left homology arm; HAR: right homology arm. (**c**) The strategy used for *Ring1b*-dTAG mouse generation. HA staining identifies the dTAG knock-in cells in blastocyst. DNA, Hoechst 33342. Scale bar, 20 μm. (**d**) Litter sizes obtained from breeding crosses between *Ring1b*^dTAG/dTAG^ females and *Ring1b*^dTAG/dTAG^ males. (**e**) Representative images of embryos collected at E7.5, E13.5 and E17.5 from *Ring1a*^dTAG/dTAG^; *Ring1b*^dTAG/+^ and *Ring1a*^dTAG/dTAG^; *Ring1b*^dTAG/+^ crosses. Asterisks indicate *Ring1a*^dTAG/dTAG^; *Ring1b*^dTAG/dTAG^ embryos. PCR-based genotyping of the corresponding embryos for *Ring1a* and *Ring1b* alleles is shown below each image set, with knock-in (KI) and wild-type (WT) bands indicated. Images are representative of at least three independent litters.

**Extended Data Figure 4.**
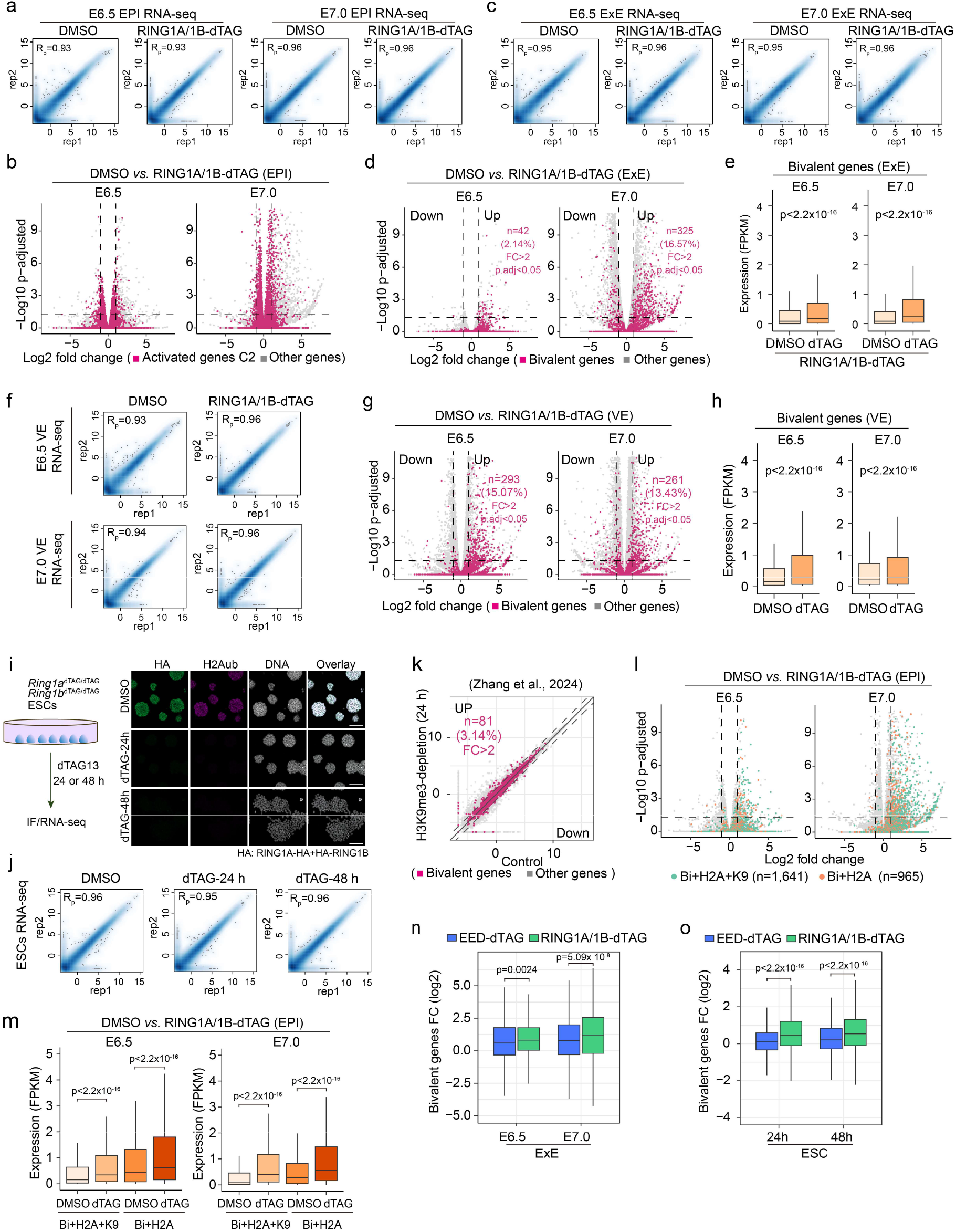
Loss of H2Aub upregulates polyvalent genes in post-implantation embryos. (**a**) Scatter plots showing the reproducibility between biological replicates for E6.5 or E7.0 EPI RNA-seq datasets. Pearson correlation coefficients are shown. (**b**) Volcano plots comparing gene expression changes between DMSO and RING1A/1B-depleted E6.5 and E7.0 EPI. Red dots represent activated genes (C2 in Fig. 2a), and gray dots represent other genes. (**c**) Scatter plots showing the reproducibility between biological replicates for E6.5 or E7.0 ExE RNA-seq datasets. Pearson correlation coefficients are shown. (**d**) Volcano plots comparing gene expression changes between DMSO and RING1A/1B-depleted E6.5 and E7.0 ExE. Red dots represent bivalent genes (n=1,961), and gray dots represent non-bivalent genes. (**e**) Box plots showing the RNA expression changes of bivalent genes in DMSO and RING1A/1B-depleted E6.5 and E7.0 ExE lineages. p values were calculated with Wilcoxon signed-rank test (two-tailed). (**f**) Scatter plots showing the reproducibility between biological replicates for E6.5 or E7.0 VE RNA-seq datasets. Pearson correlation coefficients are shown. (**g**) Volcano plots comparing gene expression changes between DMSO and RING1A/1B-depleted E6.5 and E7.0 VE. Red dots represent bivalent genes, and gray dots represent non-bivalent genes. (**h**) Box plots showing RNA expression changes of bivalent genes in DMSO and RING1A/1B-depleted E6.5 and E7.0 VE lineages. p values were calculated with Wilcoxon signed-rank test (two-tailed). (**i**) Immunostaining showing acute RING1A/1B and H2Aub degradation in ESCs following 24 and 48 h dTAG^V^-1 treatment. DNA, Hoechst 33342. Scale bar, 100 μm. (**j**) Scatter plots showing the reproducibility between biological replicates for ESCs RNA-seq datasets. Pearson correlation coefficients are shown. (**k**) Transient transcriptome sequencing (TT-seq) comparing ESCs with or without H3K9me3 depletion for 24 h (Zhang et al., 2024). (**l**) Gene expression comparison of bivalent genes with different histone markers following RING1A/1B depletion in E6.5 and E7.0 EPI. (**m**) Box plots showing RNA expression changes of bivalent genes with different histone markers in DMSO and RING1A/1B-depleted E6.5 and E7.0 EPI lineages. p values were calculated with Wilcoxon signed-rank test (two-tailed). (**n**) Comparison of log₂ fold changes of bivalent genes following EED or RING1A/1B depletion in E6.5 and E7.0 ExE. p values were calculated with Wilcoxon signed-rank test (two-tailed). (**o**) Comparison of log₂ fold changes in bivalent gene expression following EED or RING1A/1B depletion in ESCs. p values were calculated with Wilcoxon signed-rank test (two-tailed). For all the boxplots, the central band represents the median. The lower and upper edges of the box represent the first and third quartiles, respectively. The whiskers of the boxplot extend to 1.5 times interquartile range (IQR).

**Extended Data Figure 5.**
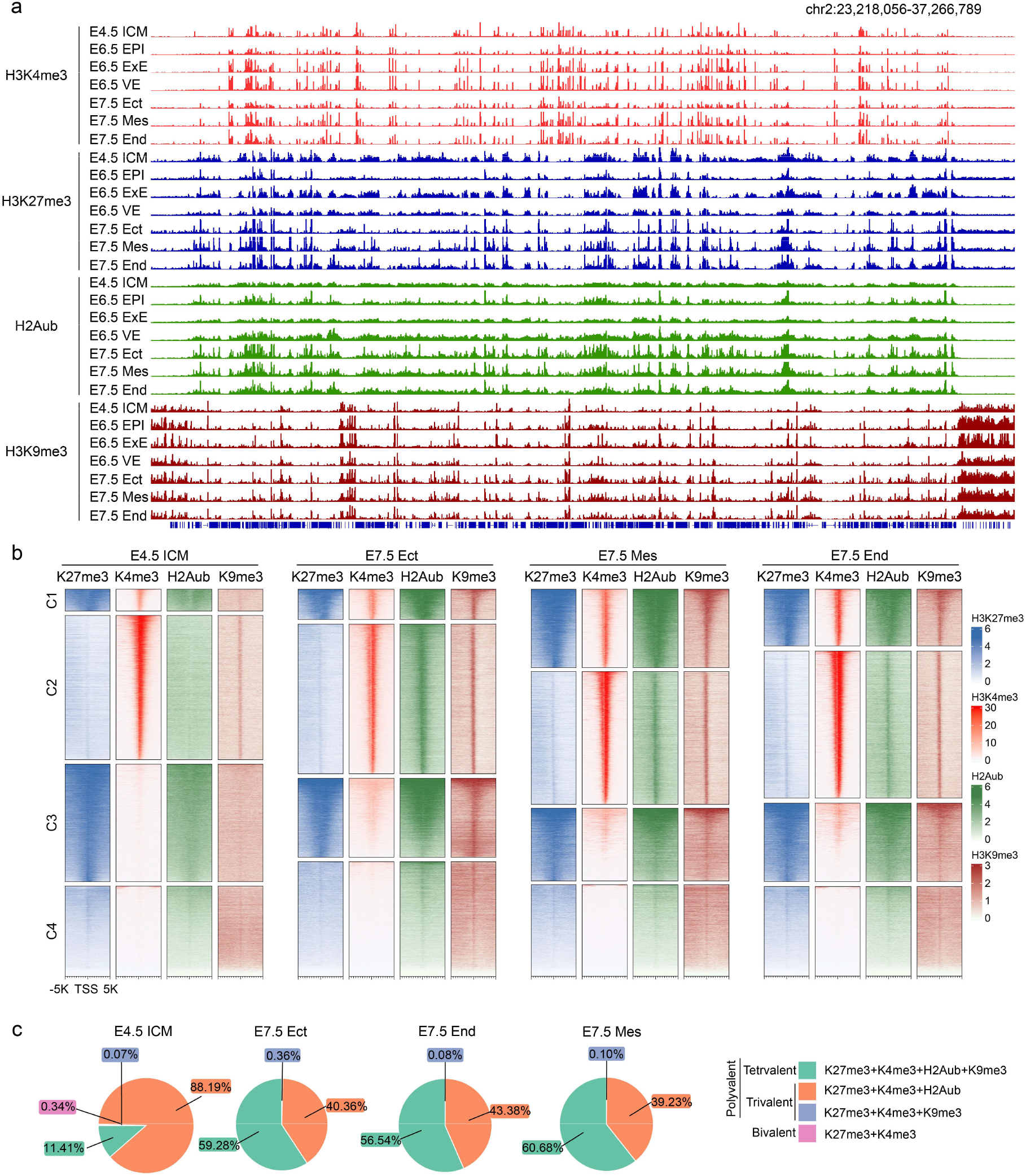
Histone modification profiling in pre- and post-implantation embryos. (**a**) Genome browser views showing examples of H3K4me3, H3K27me3, H2Aub and H3K9me3 signals in pre- and post-implantation embryos. (**b**) Histone modification profiling identifies polyvalent genes (cluster 1, C1) in E4.5 ICM, E7.5 Ect, Mes and End lineages. (**c**) Pie charts showing the proportions of genomic regions classified as bivalent, trivalent, or tetravalent chromatin states in E4.5 ICM, E7.5 Ect, End and Mes.

**Extended Data Figure 6.**
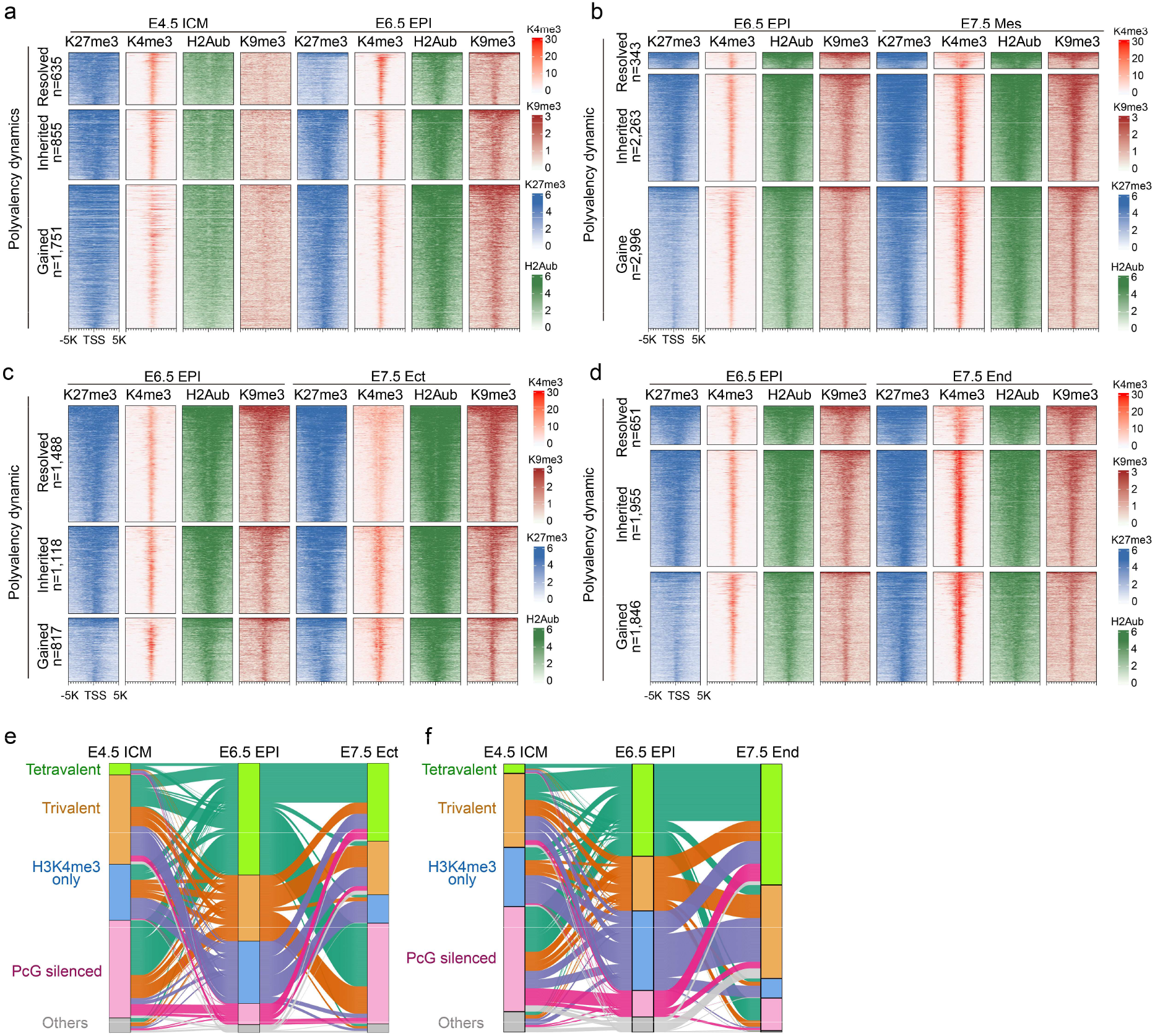
Polyvalency dynamic during development. (**a-d**) Heatmaps showing polyvalency dynamics classified as resolved, inherited or gained between consecutive developmental stages: E4.5 ICM and E6.5 EPI (**a**); E6.5 EPI and E7.5 mesoderm (Mes) (**b**); E6.5 EPI and E7.5 ectoderm (Ect) (**c**); and E6.5 EPI and E7.5 endoderm (End) (**d**). (**e, f**) Sankey diagrams illustrating transitions between chromatin polyvalency states across developmental progression from E4.5 ICM to E6.5 EPI and E7.5 Ect (**e**) or E7.5 End (**f**). Flow widths represent the proportion of genomic regions transitioning between tetravalent, trivalent, H3K4me3-only, PcG-silenced, and unmarked states. Only transitions involving tetravalent/trivalent in at least one stage were plotted.

**Extended Data Figure 7.**
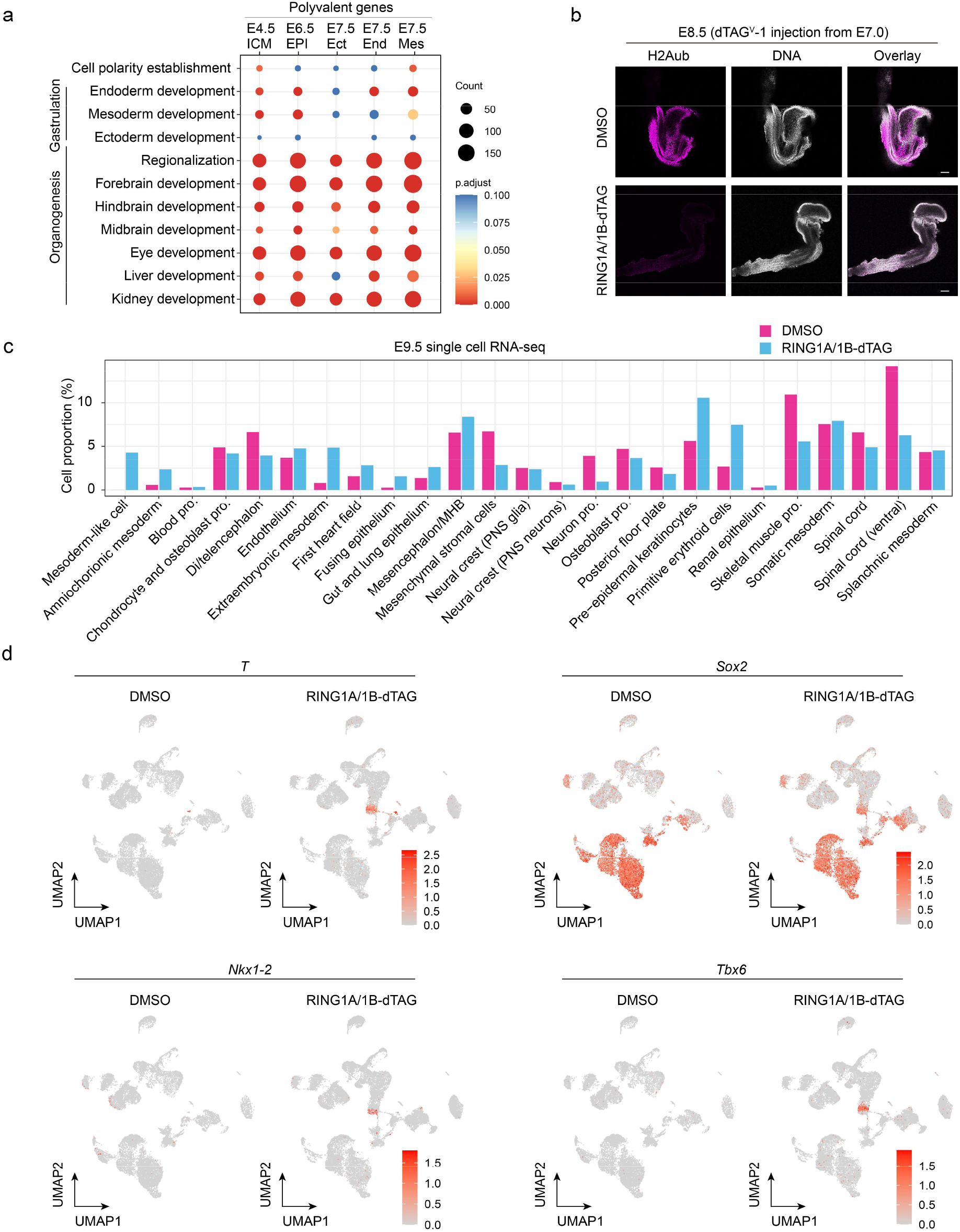
Loss of PRC1 causes E9.5 embryonic defects. (**a**) Dot plot showing Gene Ontology (GO) terms enriched among polyvalent genes in E4.5 ICM, E6.5 EPI, E7.5 Ect, End and Mes. Dot size indicates the number of genes associated with each term, and color represents the adjusted *p* value (*p.*adjust). GO terms are grouped by developmental processes related to EPI development (E6.5), gastrulation (E7.5) and organogenesis (E8.5 and later). (**b**) Immunostaining showing acute depletion of H2Aub at E8.5 following dTAG^V^-1 injection. DNA, Hoechst 33342. Scale bar, 100 μm. (**c**) Bar plot showing the proportion of cells assigned to each annotated cell type in E9.5 single-cell RNA sequencing datasets following RING1A/1B depletion. (**d**) UMAP feature plots showing expression of *T*, *Sox2*, *Nkx1-2*, and *Tbx6* in E9.5 embryos under DMSO or RING1A/1B-dTAG treatment conditions. Each dot represents a single cell, projected onto the same UMAP embedding. Color intensity indicates normalized gene expression levels.

**Extended Data Figure 8.**
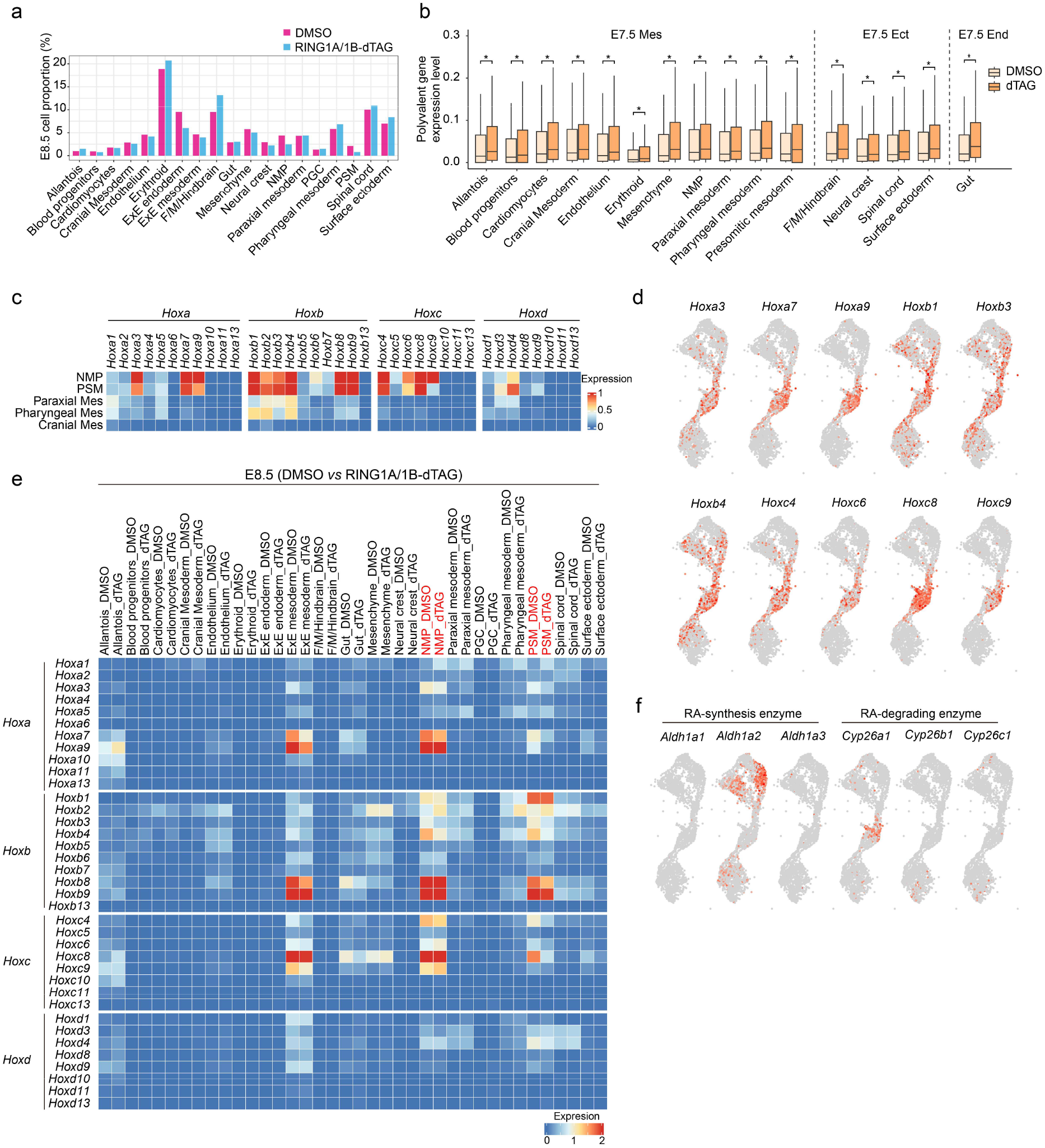
Single-cell RNA-seq analysis of E8.5 embryos with PRC1 depletion. (**a**) Proportion of cells assigned to each annotated cell type in E8.5 single-cell RNA sequencing datasets from DMSO-treated and RING1A/1B-depleted embryos. (**b**) Box plots showing expression levels of polyvalent genes across annotated cell types in E7.5 Mes, Ect, and End under DMSO or dTAG treatment. * p < 2x10^-6^. p values were calculated with Wilcoxon signed-rank test (two-tailed). In the boxplot, the central band represents the median. The lower and upper edges of the box represent the first and third quartiles, respectively. The whiskers of the boxplot extend to 1.5 times interquartile range (IQR). (**c**) Heatmap showing expression levels of *Hoxa*, *Hoxb*, *Hoxc* and *Hoxd* cluster genes across neuromesodermal progenitors (NMP), presomitic mesoderm (PSM), paraxial mesoderm, pharyngeal mesoderm and cranial mesoderm at E8.5 stage. (**d**) UMAP projections of E8.5 single-cell RNA-seq data showing the expression of selected *Hox* genes across mesoderm related cell types. (**e**) Heatmap showing expression levels of *Hoxa*, *Hoxb*, *Hoxc* and *Hoxd* cluster genes across annotated cell types of the E8.5 embryo with or without RING1A/1B depletion. (**f**) UMAP feature plots of E8.5 single-cell RNA-seq data showing expression of RA-synthesizing enzymes (*Aldh1a1*, *Aldh1a2* and *Aldh1a3*) and RA-degrading enzymes (*Cyp26a1*, *Cyp26b1* and *Cyp26c1*).

